# M-CSF drives alveolar macrophage plasticity during development and cytomegalovirus infection

**DOI:** 10.64898/2026.04.11.717936

**Authors:** Sebastian Baasch, Alina Nelipovich, Zhibek Zhumadilova, Julia Henschel, Nisreen Ghanem, Clarissa-Laura Döring, Philipp Aktories, Leonhard Wagner, Agnibesh Dey, Domien Vanneste, Martin Helmstädter, Katrin Kierdorf, Julia Kolter, Gabriele Lubatti, Sagar, Thomas Marichal, Zsolt Ruzsics, Philipp Henneke

**Affiliations:** Institute for Infection Prevention and Control, University Medical Center, Faculty of Medicine, University of Freiburg, Freiburg, Germany; Centre for Chronic Immunodeficiency, University Medical Center, Faculty of Medicine, University of Freiburg, Freiburg, Germany; Faculty of Biology, University of Freiburg, Freiburg, Germany; Department of Medicine II, Faculty of Medicine, Freiburg University Medical Center, University of Freiburg, Freiburg, Germany; Institute of Neuropathology, Medical Faculty, University of Freiburg, Freiburg, Germany; Laboratory of Immunophysiology, GIGA Institute, University of Liège, Liège, Belgium; Faculty of Veterinary Medicine, University of Liège, Liège, Belgium; Department of Medicine IV, Faculty of Medicine, Medical Center-University of Freiburg, University of Freiburg, Freiburg, Germany; EMcore, Renal Division, Department of Medicine, University Hospital Freiburg, University Faculty of Medicine, Freiburg, Germany; Centre for Integrative Biological Signalling Studies (CIBSS), University of Freiburg, Freiburg, Germany; Institute of Virology, University Medical Center, Faculty of Medicine, University of Freiburg, Freiburg, Germany

## Abstract

Alveolar macrophages (AM), the most frequent resident immune cells of the lung, are at the first line of defence against respiratory pathogens and instruct structural lung cells, *e.g.* in tissue repair. They are long-lived and receive their terminal phenotypic imprint through signals originating from the unique location at the tissue-air interface, as well as through cytokines like granulocyte-macrophage colony-stimulating factor (GM-CSF) and transforming growth factor-β (TGF-β). However, the regulatory mechanisms governing their phenotypic plasticity, which is conceptually critical for their positioning and differentiation in early life and for their functional adaptation during infection, remain poorly defined. Here we explored respiratory tract infection with cytomegalovirus (CMV), which is closely linked to mammalian immune evolution. Complementary host-pathogen fate-mapping strategies revealed AM to constitute the bottleneck for efficient mouse (M)CMV infection. MCMV infection induced macrophage colony-stimulating factor (M-CSF) in the alveolar space, and culturing of AM in M-CSF led to a profound remodelling of morphology, immunophenotype, and transcriptional identity, *e.g.* it increased the expression of interferon-stimulated genes (ISG), which modulated susceptibility to infection. Notably, already at baseline recently differentiated neonatal AM across species retained an M-CSF-associated transcriptional program. This was linked to reduced permissiveness to respiratory MCMV infection *in vivo*. Overall, our findings identify the role of M-CSF–dependent signalling in conferring plasticity to AM, when it is most needed, particularly during early-life establishment and in response to viral infection.

## Introduction

The lung comprises two main tissue compartments separated from one another by a single layer of epithelial cells. The pulmonary interstitium provides structural support and contains connective tissue, vasculature and lymphatic vessels, nerves, and airways. The alveolar space is filled with surfactant to prevent collapse of the lung and to enable gas exchange. Within this compartmentalised architecture lung macrophages segregate accordingly into alveolar macrophages (AM) and interstitial macrophages (IM). IM display greater phenotypic heterogeneity and are thought to be specialised in intercellular communication with stromal cells, endothelial cells, and lymphocytes, as well as in modulation of tissue repair and fibrosis (Chakarov et al., 2019; Gibbings et al., 2017; Li et al., 2024; Peng et al., 2025; Schyns et al., 2019; Vanneste et al., 2023). In contrast, AM adhere to the alveolar epithelium at the tissue-air interface, positioning them as the first line of defence against inhaled particles and pathogens and as key regulators of surfactant turnover (Schneider et al., 2014a; Trapnell et al., 2019).

Throughout the body, embryonic macrophage progenitors from the yolk sac rely on the macrophage colony-stimulating factor (M-CSF) receptor (Gomez Perdiguero et al., 2015). As a consecutive wave of macrophage seeding, fetal monocytes are thought to give rise to tissue resident macrophages (Ginhoux and Guilliams, 2016). Fetal monocytes have a prominent capacity to differentiate into AM in the alveolar space (Li et al., 2020; van de Laar et al., 2016). They migrate into the lung around embryonic day (E) 16.5 and differentiate through an intermediate pre-AM state into mature AM between postnatal day (PND) 1 and PND 3 (Guilliams et al., 2013). In contrast to almost all other tissue resident macrophages, including IM that rely on M-CSF (Vanneste et al., 2023), AM are established to uniquely change their dependency on M-CSF at birth and subsequently rely on the granulocyte-macrophage colony stimulating factor (GM-CSF) (Guilliams et al., 2013). GM-CSF together with transforming growth factor (TGF)-β induce the transcription factor peroxisome proliferator-activated receptor (PPAR)γ (Schneider et al., 2014b; Yu et al., 2017), which is crucial for AM tissue resident identity and function (Gautier et al., 2012). However, it remains unclear whether the developmental switch in growth factor dependence from M-CSF to GM-CSF occurs abruptly or through a gradual adaptation, and how this transition impacts AM plasticity. Moreover, the functional consequences of this perinatal shift – from M-CSF-primed macrophage progenitors to fully differentiated, GM-CSF-dependent AM – remain poorly defined.

The activation level of macrophages is highly dependent on environmental cues. For instance, during infection, pathogen-derived factors, cytokines and inflammatory signals can dynamically reprogram macrophage function, thereby shaping disease outcomes (Baasch et al., 2021; Forde et al., 2023). Given that many infections occur early in life, this adaptational state of AM during an early developmental window may critically influence immune responses and the course of disease.

Cytomegaloviruses (CMV) are ubiquitous β-herpesviruses that infect their hosts typically early in life. The lung is a primary infection site, from where the virus disseminates to draining lymph nodes and peripheral organs (Oduro et al., 2016). In immunocompetent individuals, primary infection is typically asymptomatic and leads to life-long persistence. The virus displays a broad cellular tropism with the ability to infect fibroblasts, epithelial and endothelial cells, as well as immune cell populations. Among these, myeloid cells have been implicated in early viral replication and dissemination (Baasch et al., 2020). In particular, AM have been proposed to represent an initial target population during respiratory infection (Baasch et al., 2021; Farrell et al., 2016). Nevertheless, the precise identity of the earliest infected cells in the lung and the contribution of AM to early MCMV pathogenesis remain debated. Furthermore, the cellular and molecular factors that shape the outcome of early life CMV infection of the respiratory tract remain incompletely understood.

Here, we combined *in vivo* and *ex vivo* approaches to explore AM plasticity during postnatal development and in CMV infection. We found that AM retained a high degree of phenotypic and transcriptional plasticity in response to M-CSF, which was more evident in neonatal as compared to adult AM. In MCMV infection AM were essential sites of replication. Functionally, M-CSF which was induced in MCMV infection conditioned AM toward a state of reduced susceptibility to CMV, likely mediated by elevated baseline expression of interferon-stimulated genes (ISG). Together, these findings highlight M-CSF as a key modulator of AM identity early in life and in viral infection.

## Results

### Alveolar macrophages are pivotal for early respiratory MCMV infection

AM are the initial and primary target of respiratory MCMV infection (Baasch et al., 2021; Farrell et al., 2016). They disseminate the virus locally and further to the draining lymph node (Baasch et al., 2021). However, their exact role during early MCMV pathogenesis is still under debate, in particular in relation to dendritic and epithelial cells (Farrell et al., 2019). Due to overlapping expression of CD11c with dendritic cells and rapid surface marker alterations of infected cells, we applied complementary host-pathogen fate mapping models to identify infected cells based on reporter activity. First, we infected adult cell type specific Cre-recombinase expressing mice with MCMV^LSL-GFP^ (Baasch et al., 2024, 2021; Tegtmeyer et al., 2019), in which the host Cre-enzyme removes the STOP-codon upstream of viral GFP, leading to reporter expression only in infected cells that express the recombinase (Figure S1A). Doing so, we found GFP^+^ cells only after intranasal infection of *LysM^cre/+^* and *Itgax^cre/+^* and not in *Clec9a^cre/+^* mice, suggesting that CD11c-expressing macrophages and potentially monocytes, but not dendritic cells, were productively infected early (12 hours post infection, hpi) after infection with MCMV (Figure 1A, B). In order to further specify the infected cell type, we performed thorough immunophenotyping of GFP^+^ cells before surface marker changes typically occur (Baasch et al., 2021) and identified AM to be predominantly infected (Figure 1C). Next, we expanded our cell type-specific fate mapping to all lung macrophages (*Mrc1^creER/+^*), IM (*Cx3cr1^creER/+^*) and surfactant producing alveolar epithelial cell type II (*S*ftpc*^creER/+^*). For efficient labelling we administered tamoxifen twice just before infection with MCMV*^LSL-GFP^* (Figure S1B). In accordance to the previous analyses (Figure 1 A-C), only conditional fate mapping mice, that targeted AM, expressed GFP (Figure 1D, E), which was again confirmed by immunophenotyping (Figure 1F). To avoid bias introduced by the usage of mice where only specific cell types expressed the recombinase, we infected *CAG^creER/+^* mice, which express a tamoxifen-inducible creER driven by a CMV-IE enhancer and chicken β-actin / rabbit β-globin hybrid promoter in all cells (Figure S1B). Doing so, we confirmed that even if the recombinase is expressed ubiquitously, AM are the dominant cell type harbouring MCMV (Figure 1G-I). To challenge the findings obtained with a MCMV-associated reporter strain, we infected *R26*^LSL-Tomato^ mice with a newly developed MCMV*^cre^*. To test functional cleavage via viral recombinase production, we generated bone marrow-derived macrophages (BMDM) from *R26^LSL-Tomato/+^* mice, infected these with a multiplicity of infection (MOI) 5 of MCMV*^cre^* and monitored reporter expression. We observed surface marker downregulation of CD45 and F4/80 on infected BMDM 3 days post infection (dpi), as reported before (Figure S1C; (Baasch et al., 2021)). While almost all BMDM were infected using MOI 5 already at 1 dpi, the intensity of reporter expression steadily increased over time (Figure S1D). Given that Tomato expression increases over time (Figure S1D), we extended the time point of analysis (Figure S1E) and readily detected infected cells at 36 hpi, independent of whether mice carried the heterozygous or homozygous reporter allele for *R26^LSL-Tomato/+^* (Figure 1J, K). The majority of Tomato^+^ infected cells were again identified as AM (Figure 1L).

**Figure 1:**
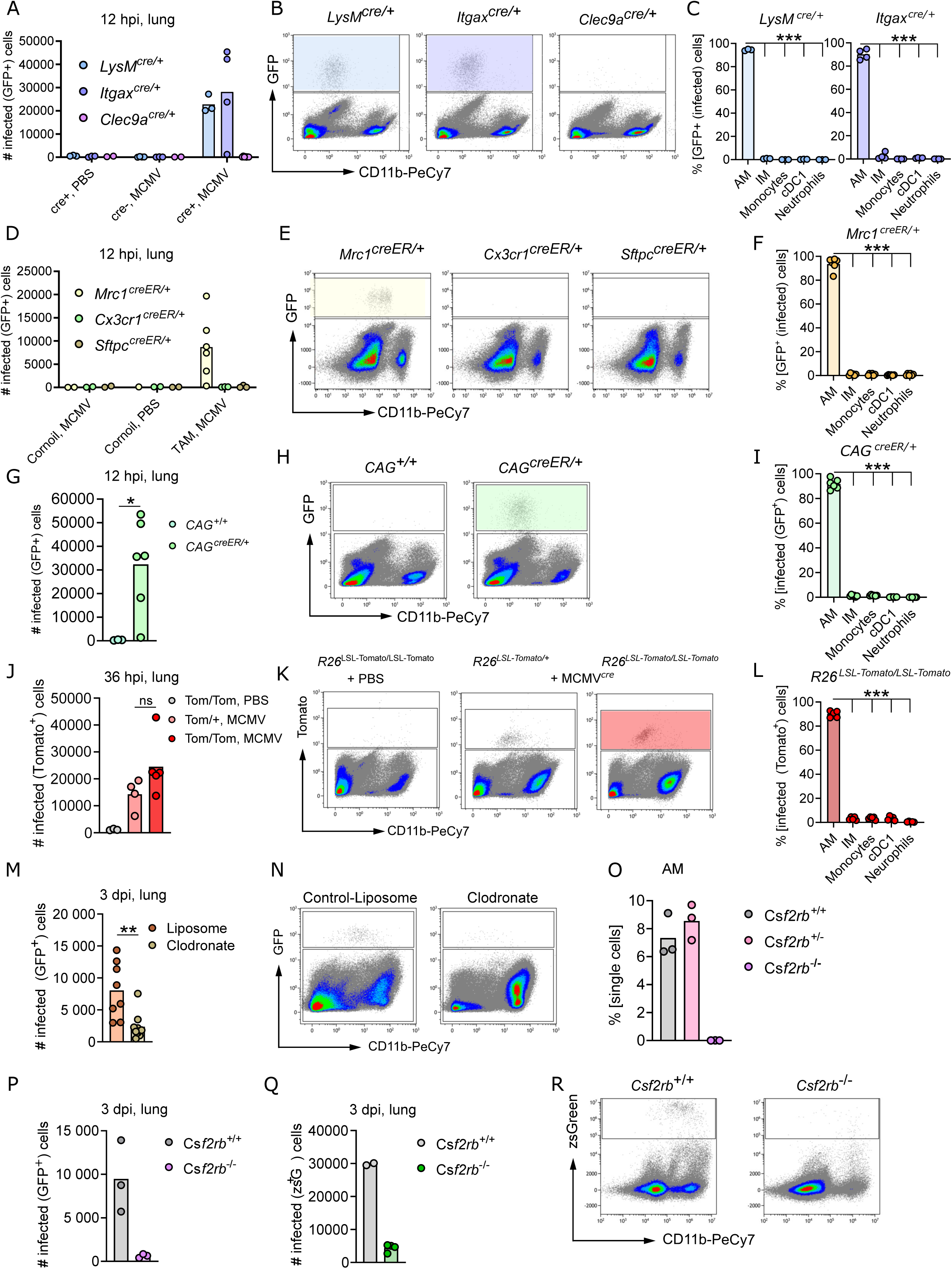
MCMV predominantly infects AM during initial infection (A) Quantification infected GFP^+^ cells after intranasal infection of *LysM*^cre/+^ *Itgax*^cre/+^ and *Clec9a^cre/+^* mice with MCMV*^LSL-GFP^* at 1 dpi. (B) Representative flow cytometry plots of MCMV*^LSL-GFP^* infected lungs (A), pregated from single living cells. (C) Quantification of GFP^+^ infected cell types. Cell types were defined as described in (Baasch et al., 2024). One-way ANOVA. (D) Quantification of infected GFP^+^ cells after intranasal infection of TAM-induced *Mrc1^creER/+^, Cx3cr1^creER/+^* and Sftpc*^creER/+^* mice with MCMV*^LSL-GFP^* at 1 dpi. (E) Representative flow cytometry plots of MCMV*^LSL-GFP^* infected lungs (E), pregated from single living cells. (F) Quantification of GFP^+^ infected cell types. Cell types were defined as described in (Baasch et al., 2024). One-way ANOVA. (G) Quantification of infected GFP^+^ cells after intranasal infection of TAM-induced *CAG^creER/+^* mice with MCMV*^LSL-GFP^* at 1 dpi. Mann-Whitney test. (H) Representative flow cytometry plots of MCMV*^LSL-GFP^* infected lungs (G), pregated from single living cells. (I) Quantification of GFP^+^ infected cell types. Cell types were defined as described in (Baasch et al., 2024). One-way ANOVA. (J) Quantification of infected Tomato^+^ cells after intranasal infection of *R26^LSL-Tomato/+^* and *R26^LSL-Tomato/LSL-Tomato^* mice with MCMVLSL-GFP at 1 dpi. (K) Representative flow cytometry plots of MCMV*^LSL-GFP^* infected lungs (J), pregated from single living cells. (L) Quantification of Tomato^+^ infected cell types. One-way ANOVA. (M) Quantification of GFP^+^ infected cells after depletion of AM with clodronate liposomes (Figure S1G). Mann-Whitney test. (N) Representative flow cytometry plots of MCMV*^LSL-GFP^* infected lungs after AM depletion with clodronate liposomes or administration with control liposomes, pregated from single living cells. (O) Flow cytometric quantification of AM in *Csf2rb2*^+/+^, *Csf2rb2*^+/-^ and *Csf2rb2*^-/-^ mice. (P) Quantification of GFP^+^ infected cells after intranasal infection of *Csf2rb2*^+/+^ and *Csf2rb2*^-/-^ mice with MCMV*^GFP^*, 3 dpi. (Q) Quantification of zsGreen^+^ infected cells after intranasal infection of *Csf2rb2*^+/+^ and *Csf2rb2*^-/-^ mice with MCMV*^zsG^*, 3 dpi. (R) Representative flow cytometry plots of MCMV*^zsG^* infected lungs (Q), pregated from single living cells. Quantified data is presented as individual mice, bars indicate mean. \**p <*0.05, \*\**p <*0.01, \*\*\**p <*0.001

Together, using a variety of host-pathogen fate mapping approaches, we confirmed that AM are the predominant infected cells early after intranasal MCMV infection.

However, MCMV has a broad cellular tropism and the capacity to infect the majority of cell types in the body, ranging from fibroblasts to endothelial cells and myeloid immune cells. Therefore, we wanted to investigate the course of acute MCMV infection in the absence of AM, i.e. in pharmaceutical and genetic depletion models. First, we administered clodronate liposomes intranasally, which substantially reduced AM, and subsequently infected the mice with MCMV*^GFP^* (Figure S1F, G). Despite the broad cellular tropism, the reduction of AM also led to a diminished number of infected GFP^+^ cells (Figure 1M, N). Because, temporal depletion of macrophages, for example via clodronate liposomes, leads to rapid monocyte influx and differentiation, we further infected *Csf2rb*^-/-^ mice that lack the receptor for GM-CSF and are void of AM (Figure 1O, (Guilliams et al., 2013)) and again we found almost no infected cells 3 dpi (Figure 1P). To exclude that m36 driven GFP expression in MCMV*^GFP^* (Baasch et al., 2021) was specific for macrophages (Daley-Bauer et al., 2017; Ebermann et al., 2012) and, although expressed in other MCMV-infected cells, would favour detection of infected macrophages, we generated a reporter MCMV, which expresses zsGreen (MCMV*^zsG^*) under the control of the strong *hIE1* promotor – similar to the MCMV*^LSL-GFP^*. Notably, the capacity of initial infection and dissemination in MCMV*^zsG^* was comparable to the MCMV wild type (wt) strains Smith and Rep3.3 (Jordan et al., 2011) (Figure S1H). Intranasal infection with MCMV*^zsG^* also led to a strong decrease in overall infected cells in the absence of AM in *Csf2rb*^-/-^ mice (Figure 1Q, R).

In summary, AM are the non-redundant initial targets of MCMV replication in respiratory tract infection.

### *Csf1r* deficiency reduces AM numbers at steady state

Although CMV infections usually do not cause overt symptoms in immunocompetent hosts, the virus engages a variety of pathogen recognition receptors and induces inflammation (Baasch et al., 2020). To define the landscape of immune mediators in the lung after MCMV infection, we performed multiplex cytokine analysis of bronchoalveolar lavage fluid (BALF) at 3 dpi. As expected, we detected elevated levels of inflammatory cytokines, such as tumor necrosis factor (TNF)-α, Interleukin (IL)-6, interferon (IFN)-α and IL-1β (Figure 2A). In notable addition to these inflammatory mediators, we also observed highly elevated levels of M-CSF (Figure 2A). AM are established to receive identity-defining programming by GM-CSF and TGF-β, which induce the lineage defining transcription factor peroxisome proliferator-activated receptor (PPAR) γ (Schneider et al., 2014b). However, the impact of high M-CSF concentration, as apparently present in the alveolar space during infection and not during steady state, on adult mature AM was unclear. Accordingly, we analysed a previously published bulk RNA-sequencing dataset of control, non-infected bystander and infected AM (Baasch et al., 2021). Whereas infected cells were significantly reduced in *Csf1r* expression, control and bystander AM expressed comparable levels, suggesting the ability of uninfected AM to receive M-CSF receptor downstream signals (Figure 2B). CD169 in the lung is expressed on AM and a small subset of nerve-associated interstitial macrophages (Ural et al., 2020). Therefore, we used *Cd169^cre/+^;Csf1r^fl/fl^* mice to deplete the M-CSF receptor on AM and subsequently infected them with MCMV*^zsG^*. Under these conditions the number of infected AM was reduced at 3 dpi (Figure 2C). Notably, deletion of *Csf1r* decreased the proportion and number of AM in conditional *Cd169^cre/+^;Csf1r^fl/fl^* mice and in *Csf1r^ΔFIRE/ΔFIRE^* mice with an ubiquitous knock out of the *fms*-intronic regulatory element (FIRE) enhancer (Rojo et al., 2019) already under steady state conditions (Figure 2D-E, S2A-B), while IM were not affected (Figure S2C).

**Figure 2:**
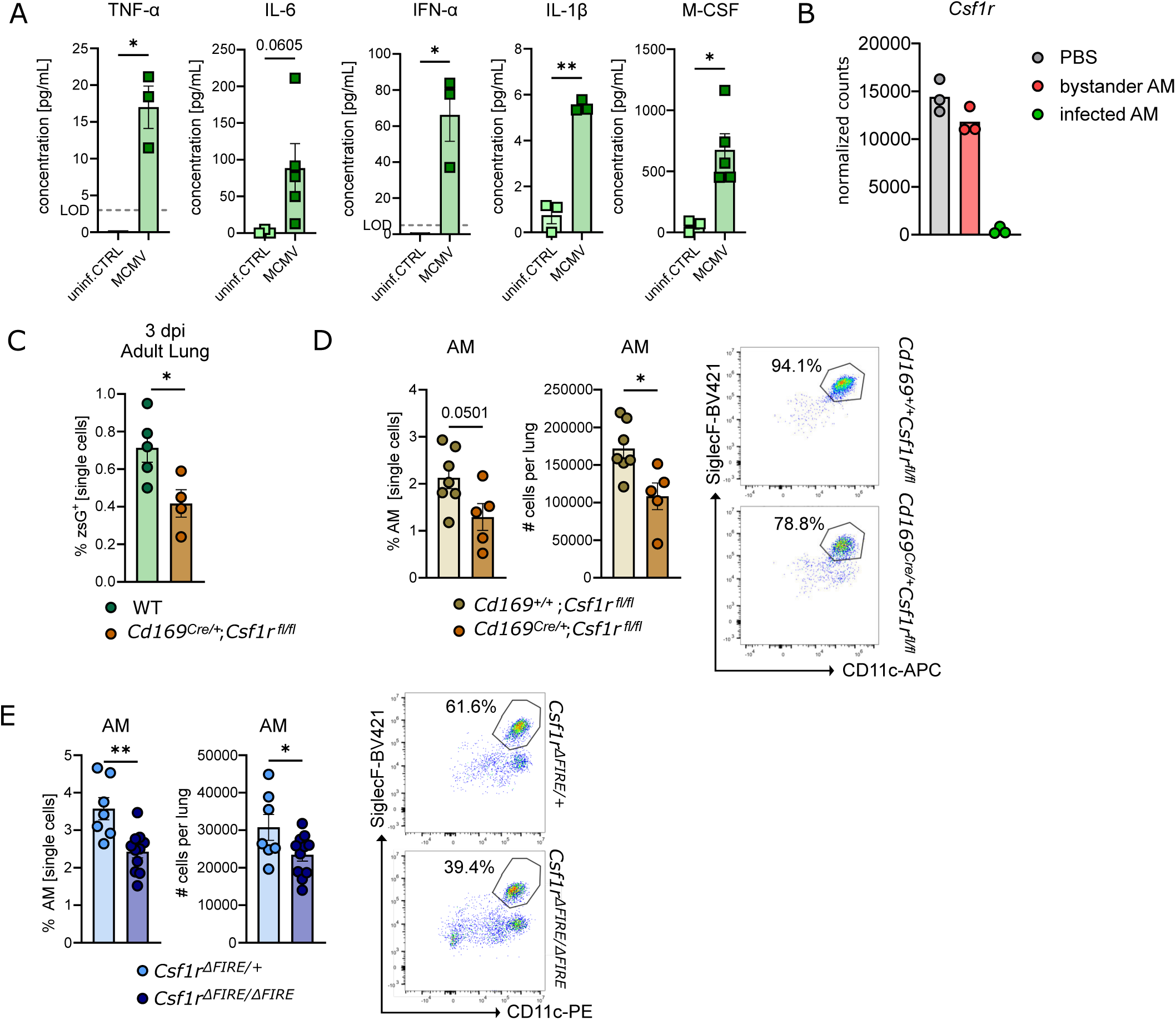
C*s*f1r deficiency reduces AM numbers at steady state. (A) Cytokine concentrations in the bronchoalveolar lavage fluid (BALF) of adult WT mice infected with MCMV*^zsG^* (1×10^5^ PFU *i.n.*) at 3 dpi or uninfected control, measured by LEGENDplex^TM^. *n=* 3-5, 2-3 independent experiments, LOD=limit of detection, unpaired t-test with Welch’s correction and unpaired t-test (M-CSF) (B) Comparison of normalised counts of *Csf1r* expression on non-infected, bystander and infected AM 3 dpi from a previously published bulkRNAseq data set (Baasch *et al* (2021), GEO accession number GSE 159389). *n=*3 (C) Percentages of zsGreen^+^ cells at 3 dpi in adult lung of WT mice in comparison to *Cd169^Cre/+^Csf1r^fl/fl^* mice infected with MCMV*^zsG^* (1×10^5^ PFU *i.n.*), assessed by flow cytometry. *n=*4-5, 4 independent experiments, unpaired t-test (D) Quantification of absolute AM numbers and percentages in *Cd169^Cre/+^Csf1r^fl/fl^* mice in comparison to *Cd169^+/+^Csf1r^fl/fl^* mice assessed by flow cytometry (left) with exemplary dot plots showing the AM population, pre-gated on living CD45^+^ Lin (CD103, CD3e, Ly6G, Ly6C)^-^ CD206^+^ cells (right). *n=*5-7, 3 independent experiments, unpaired t-test (E) Quantification of absolute AM numbers and percentages in *Csf1r^ΔFIRE/ΔFIRE^* mice in comparison to *Csf1r^ΔFIRE/+^* mice assessed by flow cytometry (left) with exemplary dot plots showing the AM population, pre-gated on living CD45^+^ Lin (CD103, CD3e, Ly6G, Ly6C)^-^ CD206^+^ cells (right). *n=*7-11, 4 independent experiments, unpaired t-test

Altogether, MCMV infection leads to high levels of M-CSF in the alveolar space and loss of M-CSF receptor expression in macrophages reduces the AM number in homeostasis and thus the target cells for MCMV infection.

### AM respond to M-CSF with morphologic, immunophenotypic and transcriptomic changes and decreased susceptibility to MCMV *ex vivo*

To assess the effects of M-CSF on adult AM in more detail, we cultured fluorescence-activated cell sorted AM either in medium that comprised GM-CSF, TGF-β and the anti-diabetic drug rosiglitazone to promote *Pparγ* expression (3x AM, (Gorki et al.; Luo et al., 2021)) or in M-CSF containing medium (M-CSF AM) (Figure S3A), and performed phenotypic and functional assays. Using scanning electron microscopy, we found that 3x AM displayed a round morphology with thin protrusions, while the majority of M-CSF cultured AM spread out and occupied a larger surface area after 9 days in culture (Figure 3A). Immunofluorescence microscopy confirmed the morphological changes in cultured M-CSF AM and revealed further surface marker changes (Figure 3B, C, S3B). 3x AM maintained characteristic surface markers SiglecF, CD11c and lacked surface expression of CX3CR1 and CD11b in microscopy, as well as flow cytometry (Figure 3B-D). However, M-CSF AM downregulated SiglecF and CD11c *ex vivo*, while upregulating CX3CR1 and CD11b over time (Figure 3B-D). To comprehensively depict the broad changes induced by M-CSF in AM we performed bulk RNA-sequencing of cells that were cultured for 9 days in either 3x or M-CSF conditions and included freshly isolated primary AM that rely on endogenous GM-CSF (Guilliams et al., 2013) and IM that rely on M-CSF (Vanneste et al., 2023) in our analysis (Figure S3C). While 3x AM retained a transcriptomic signature similar to freshly sorted AM (Figure 3F-G, Figure S3D), M-CSF AM were distinct from both, 3x cultured and bona fide AM (Figure 3E-G). Analysis of the top 100 highly variable genes and hierarchical clustering confirmed overlapping gene expression profiles of primary AM and 3x AM, as well as IM and M-CSF AM (Figure 3F). M-CSF receptor signalling activates the transcription factor MAFB, which in turn regulates expression of core macrophage genes including those encoding for C1q complement members (*C1qa*, *C1qb*, *C1qc*), folate receptor beta (*Folr2*), apolipoprotein E (*Apoe*) and M-CSF receptor (*Csf1r*) (Vanneste et al., 2026). Accordingly, genes, such as *Mafb*, *C1qa, C1qb, C1qc, Folr2, Apoe* and *Cx3cr1* were highly transcribed in M-CSF dependent primary IM and M-CSF AM (Figure 3E-G), while primary AM and 3x AM expressed hallmark AM transcripts, such as *Ear1*, *Car4* and *Fabp1* (Gautier et al., 2012) (Figure 3E-G). To confirm the role of the M-CSF-MAFB-pathway, we applied a *Mafb* signature score (Vanneste et al., 2026) to our sequencing data and saw higher expression of MAFB induced genes in IM and M-CSF AM (Figure S3E). AM have a high capacity for phagocytosis and constantly take up surfactant and cellular debris. At the same time, they are primary targets for MCMV infection (Figure 1). Therefore, we tested the phagocytic capacity and viral susceptibility of 3x AM and M-CSF AM. First, we incubated them with fluorescent latex beads and observed a higher uptake of particles in 3x AM as compared to M-CSF AM (Figure 3H). Next, we infected cultured AM *ex vivo* with MCMV*^zsG^* (MOI1) and observed a decreased susceptibility to MCMV infection in M-CSF AM (Figure 3I). Interestingly, this resistance to viral infection was associated with a higher base line expression of ISG in M-CSF AM (Figure 3J).

**Figure 3:**
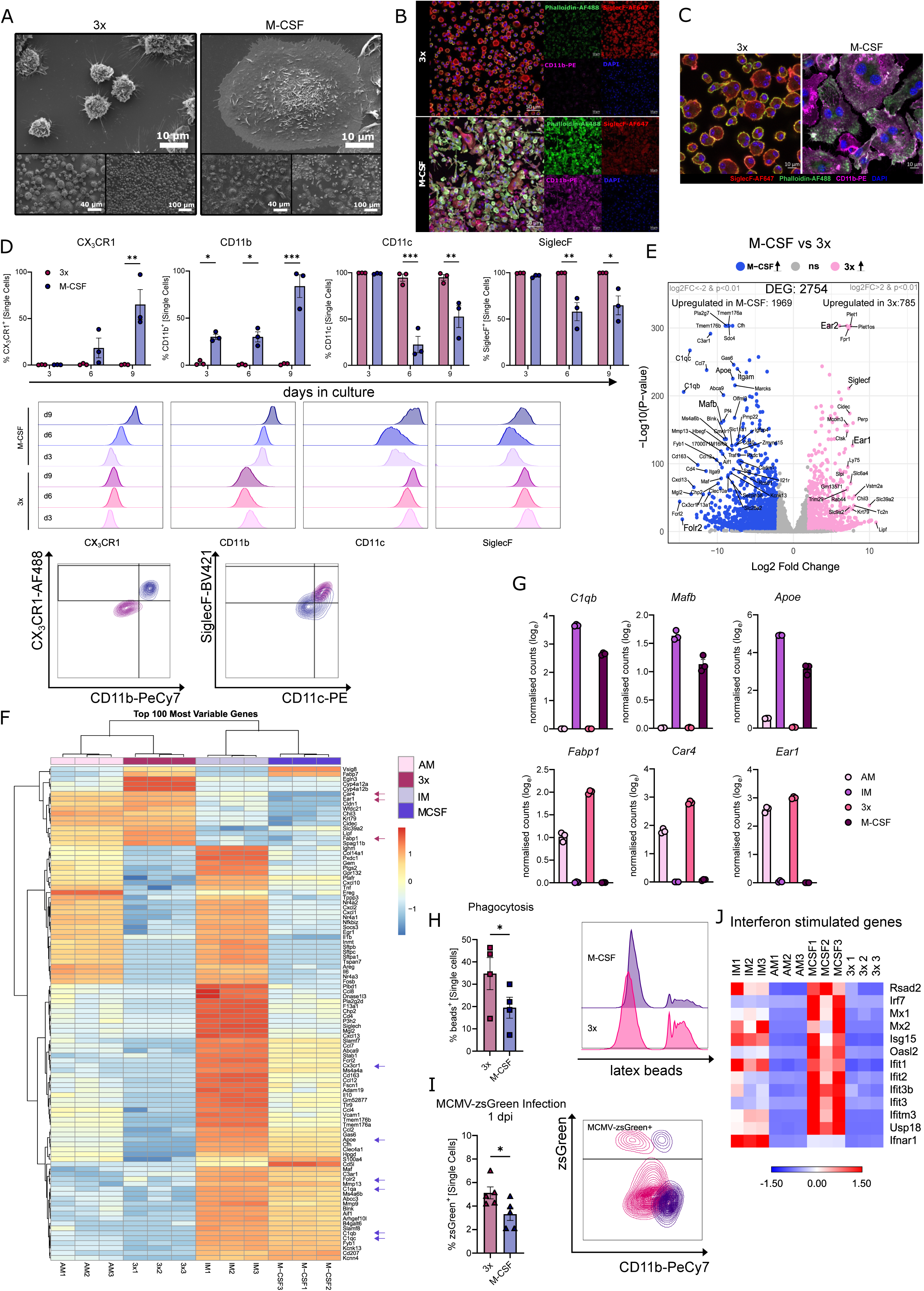
AM respond to M-CSF with morphologic, immunophenotypic and transcriptomic changes and decreased susceptibility to MCMV *ex vivo* (A) Scanning electron microscopy pictures of 3x and M-CSF cultured AM, 6500x, 800x and 350x magnifications, scale bar size is indicated in the respective pictures (B) Representative confocal images of 3x and M-CSF cultured primary mouse AM on day 9 (d9) of culture, 20x magnification, scale bar indicative of 50 µm (C) Representative pictures of cell morphology of 3x and M-CSF cultured AM on d9 of culture, 63x magnification, scale bar indicative of 10 µm (D) Flow cytometrically assessed percentages of CX3CR1^+^, CD11b^+^, CD11c^+^, and SiglecF^+^ primary mouse AM, cultured in either 3x or M-CSF, at d3, d6 and d9 of culture (top) and exemplary histograms of the respective markers across different timepoints together with a representative contour plot of 3x and M-CSF cultured cell populations at d9 of culture (bottom). *n=*3, 3 independent experiments, Tukey’s multiple comparisons test (E) Volcano plot showing differentially expressed genes (coloured dots: abs(log2_FC) > 2 and *p* < 0.01, labelled dots: abs(log2-FC) > 7, *p* < 0.01) that are upregulated in 3x (rosa) or upregulated (purple) in M-CSF cultured AM from *Cx3cr1*^gfp/+^ reporter mice. *n=*3 (F) Heatmap depicting the 100 most variable genes between 3x cultured AM, M-CSF cultured AM, as well as freshly sorted AM and IM from lung tissue, resulting from bulk RNA-sequencing of *Cx3cr1*^gfp/+^ reporter mice. (G) Selected differentially expressed genes from bulkRNAseq of 3x and M-CSF cultured AM from *Cx3cr1^gfp/+^* reporter mice alongside freshly sorted AM and IM as controls. *n=*3 (H) Assessment of phagocytosis in 3x and M-CSF cultured AM with yellow-green fluorescent latex-beads using flow cytometry at d9 of culture (left) and appropriate representative histogram (right). *n=*4, 4 independent experiments, paired t-test (I) Quantification of MCMV*^zsG^* (MOI 1) infected 3x and M-CSF cultured AM at 1 dpi (left) and respective representative contour plots (right). n=5, 4 independent experiments, paired t-test (J) Heatmap, obtained from bulkRNAseq, illustrating interferon-stimulated genes (ISGs) in 3x and M-CSF culture AM on d9 of culture, as well as freshly isolated AM and IM. Data are represented as mean ± SEM. \**p <*0.05, \*\**p <*0.01, \*\*\**p <*0.001

Overall, M-CSF induced a change in morphology, surface marker expression and gene signature in AM *ex vivo.* Moreover, M-CSF increased expression of ISG in AM that was associated with a reduced susceptibility to MCMV infection.

### Neonatal AM arise from M-CSF experienced progenitors and retain a M-CSF signature postnatally

AM were shown to originate from fetal monocytes that seed the lung around E16.5. In mice, they transition through a pre-AM stage shortly after birth and mature into SiglecF^+^ CD11c^+^ cells within the first week of life (Guilliams et al., 2013). Embryonic progenitors in the yolk sac express *Csf1r* (Gomez Perdiguero et al., 2015) and disruption of M-CSF signalling reduces the earliest macrophage progenitors during embryogenesis (Ginhoux et al., 2010; Hoeffel et al., 2012), as well as tissue-resident macrophages in the adult mouse (Dai et al., 2002; Hoeffel et al., 2012; Kolter et al., 2019; MacDonald et al., 2010). These observations suggest that the intermediate progenitors, i.e., fetal monocytes and macrophage progenitors, are also exposed to M-CSF signalling during development. We therefore hypothesized that embryonic precursors of AM display a strong M-CSF associated transcriptional program and that they subsequently transition to overall dependence on GM-CSF signalling. To test this hypothesis, we analysed monocytes and fetal macrophages from the lung, pre-AM and mature AM around birth (E18.5- 3 PND, (Guilliams et al., 2013)). Consistent with ongoing differentiation, the proportion of fetal macrophages and pre-AM decreased after birth, whereas monocyte and mature AM populations progressively increased (Figure 4A, B). To define the transcriptional programs associated with these developmental stages, we sorted these cell populations at PND 3 together with mature AM from adult mice to perform bulk RNA-sequencing. Hierarchical clustering of the top 1,000 differentially expressed genes revealed three gene clusters (Figure S4A). Clusters 1 and 3 consisted of genes that increased from pre-AM to adult AM, whereas cluster 2 consisted of genes that decreased from pre-AM to adult AM. Next, we used our previously described *ex vivo* bulk RNA sequencing data (Figure 3) to derive a set of M-CSF dependent and AM signature genes. Specifically, we compared IM and M-CSF AM, as well as primary AM and 3x AM, and defined intersecting transcripts as M-CSF- or AM-associated gene sets, respectively (Figure S4B). Subsequently, we tested which clusters (Figure S4A) showed enrichment for the defined M-CSF or AM signatures. Clusters 1 and 3 showed enrichment for the AM signature (Figure 4C), while cluster 2 showed enrichment for the M-CSF signature (Figure 4D). In this approach, adult AM, as expected, showed the strongest AM signature, expressing markers like *Ear1*, *Car4* and *Fabp1*, while fetal macrophages and pre-AM showed a less pronounced AM signature (Figure 4C, E). Neonatal AM also showed overall reduced expression of AM specific genes (Figure 4C), including *Ear1* and *Fabp1*, but with the exception of *Car4* (Figure 4E). M-CSF dependent genes, i.e., *Mafb*, *C1qb* and *Apoe*, were strongly upregulated in pre-AM and fetal macrophages (Figure 4D-E). Additionally, neonatal AM, in contrast to adult AM, still showed higher expression of M-CSF dependent genes, such as *Mafb* and *Apoe* (Figure 4D-E). These results suggest that at PND 3, mature neonatal AM, defined as SiglecF⁺ CD11c⁺, incorporate signals from new environmental cues, leading to upregulation of AM signature genes, while retaining a residual M-CSF-associated transcriptional imprint as apparent by increased *Mafb* and *Apoe* expression. Principal component analysis (PCA) confirmed the transcriptional difference between neonatal and adult AM (Figure S4C).

**Figure 4:**
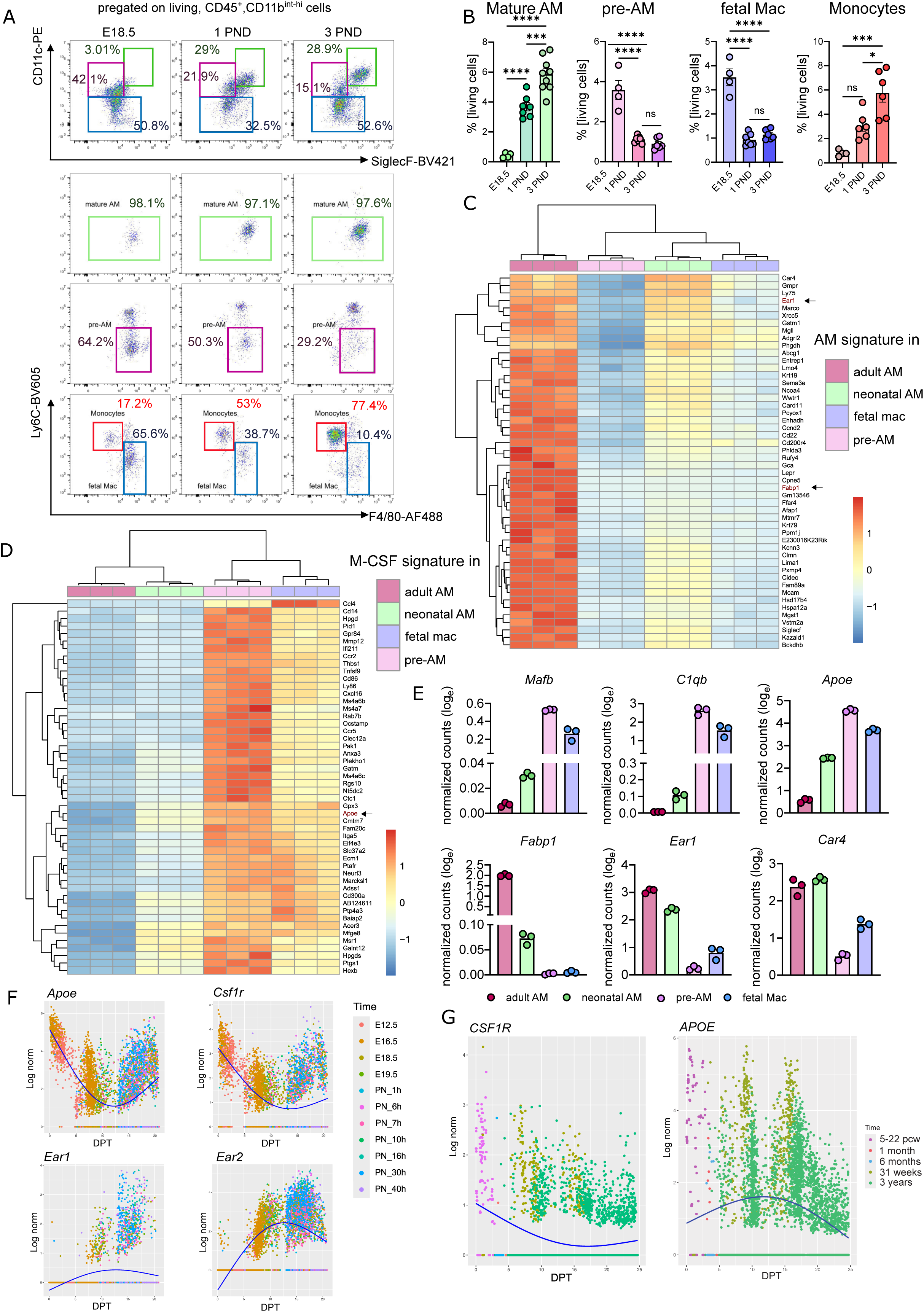
Neonatal AM arise from M-CSF experienced progenitors and retain a M-CSF signature postnatally (A) Gating strategy for mature AM, pre-AM, fetal macrophages (fetal mac) and monocytes from E18.5, 1 PND and 3 PND WT mice (adapted from Guilliams *et al* 2013). (B) Quantification of living mature AM, pre-AM, fetal macrophages and monocytes, assessed by flow cytometry at E18.5, 1 PND and 3 PND. *n=*4-6, 1-2 independent experiments, one-way ANOVA (C) Heatmap of mouse bulk RNA-sequencing showing top 50 genes that are common between the AM signature (Figure S4B) and the clusters 1 and 3 (= clusters including genes with increasing expression from pre-AM to adult AM condition, Supplemental Figure 4A). *n=*3 (D) Heatmap of mouse bulk RNA-sequencing showing top 50 genes that are common between the M-CSF signature (Figure S4B) and the cluster 2 (= cluster including genes with increasing expression from AM and to pre-AM, Supplemental Figure 4A). *n=*3 (E) Selection of normalised counts in bulkRNA sequenced adult AM, neonatal AM, pre-AM and fetal Mac for exemplary M-CSF signature genes (*Mafb, C1qb, Apoe*) and AM signature genes (*Fabp1, Ear1*). *n=*3 (F) Expression dynamics of *Apoe, Csf1r, Ear1* and *Ear2* along the diffusion pseudo time in mouse scRNA-sequencing data (Cohen et al., 2018). Fitted line is obtained with Generalised Additive Model (GAM) fit. (G) Expression dynamics of *APOE* and *CSF1R* along the diffusion pseudotime in human data (Barnes et al., 2023; Sikkema et al., 2022; Wang et al., 2020). Fitted line is obtained with Generalised Additive Model (GAM) fit. Data are represented as mean ± SEM. \**p <*0.05, \*\*\**p <*0.001, \*\*\*\**p <*0.0001, *ns=* non-significant.

To validate these findings in an independent dataset, we reanalysed a published single cell RNA sequencing dataset of the developing mouse lung spanning from prenatal to postnatal time points (Cohen et al., 2018). Macrophages expressing *Apoe* and *Csf1r* were predominantly detected at E12.5 and E16.5 and transitioned towards mature AM until 40 hours after birth, as inferred by pseudotime analysis and gradual expression of canonical AM markers *Ear1* and *Ear2* (Figure 4F). Using another publicly available single cell RNA-sequencing datasets from the fetal (5-22 post conceptional weeks) and postnatal lung (1, 6 months and 3 years postnatal)(Barnes et al., 2023; He et al., 2022; Sikkema et al., 2022; Wang et al., 2020), we sought to verify our findings during human development. Similarly to our findings in mice, human developmental data revealed a trajectory in which early *CSF1R*^+^ *APOE*^+^ monocyte/macrophages transitioned to mature AM (Figure 4G).

To determine whether this developmental trajectory corresponded to changes in M-CSF-associated transcriptional signatures, i.e. higher at earlier time points on the progenitor level and lower at later time points, when mature AM are present, we first built a trajectory of cells and subsequently assigned a diffusion pseudotime (DPT) to each cell, which correlated with the real time points. We then analysed gene expression dynamics across developmental trajectories by clustering transcripts with similar temporal expression patterns along the DPT. This approach identified four gene clusters in the human datasets and six clusters in the mouse dataset (Figure S4D-E). In human datasets, cluster 1 genes that progressively increased over developmental time were significantly enriched for transcripts associated with the AM signature (Figure S4D, F). In contrast, clusters 3 and 4, which showed an overall decrease in gene expression during development, were significantly enriched for transcripts associated with the M-CSF signature (Figure S4D, F). A similar pattern emerged in the mouse dataset: clusters 5 and 6, characterized by overall increasing gene expression over time, were enriched for AM-associated genes, whereas clusters 2 and 4, which displayed declining expression during development, were enriched for M-CSF–associated transcripts (Figure S4E, G).

Together, these analyses revealed a conserved developmental trajectory in which early macrophage progenitors displayed a transcriptional program associated with M-CSF signalling that progressively diminished as cells differentiated into mature AM. Furthermore, the findings suggest that developmental exposure to M-CSF leaves a transient molecular imprint during AM maturation that is gradually replaced by GM-

CSF-dependent transcriptional programs after birth, but is still in part present in neonatal mature AM.

### Neonatal mice are less susceptible to respiratory MCMV infections

Finding that early after birth, an M-CSF signature was still present in AM, we next investigated whether neonatal AM showed higher plasticity when cultured with M-CSF. Indeed, neonatal AM downregulated SiglecF and upregulated CD11b already 3 days and 6 days after culturing in M-CSF, respectively, and thus, faster as compared to adult M-CSF AM (Figure 5A, B). CX3CR1 upregulation in neonatal M-CSF AM appeared stronger than in adult M-CSF AM, however, this was not statistical significant (Figure 5A). Functionally, neonatal 3x AM showed a similar phagocytic capacity as compared to adult M-CSF AM (Figure 5C), indicative of a partial M-CSF imprint after birth. Nonetheless, the phagocytic capacity was further decreased in neonatal M-CSF AM in comparison to neonatal 3x AM (Figure 5C). Next, we tested the susceptibility to MCMV and found that neonatal 3x AM were even less infectable than adult M-CSF AM, while neonatal M-CSF AM showed the least infectibility with MCMV at 1 dpi (Figure 5D). These findings are in line with gene ontology overrepresentation analysis of biological processes from the bulk RNA-sequencing data of mature adult and neonatal AM (Figure 4), which revealed distinct functional programs (Figure 5E). Neonatal AM were enriched for genes involved in the regulation of IFN responses, suggesting a reduced susceptibility to viral infection (Figure 5E). In contrast, adult AM showed enrichment of processes related to fatty acid metabolism and responses to oxygen-containing compounds, as well as macrophage activation and responses to bacterial stimuli (Figure 5E).

**Figure 5:**
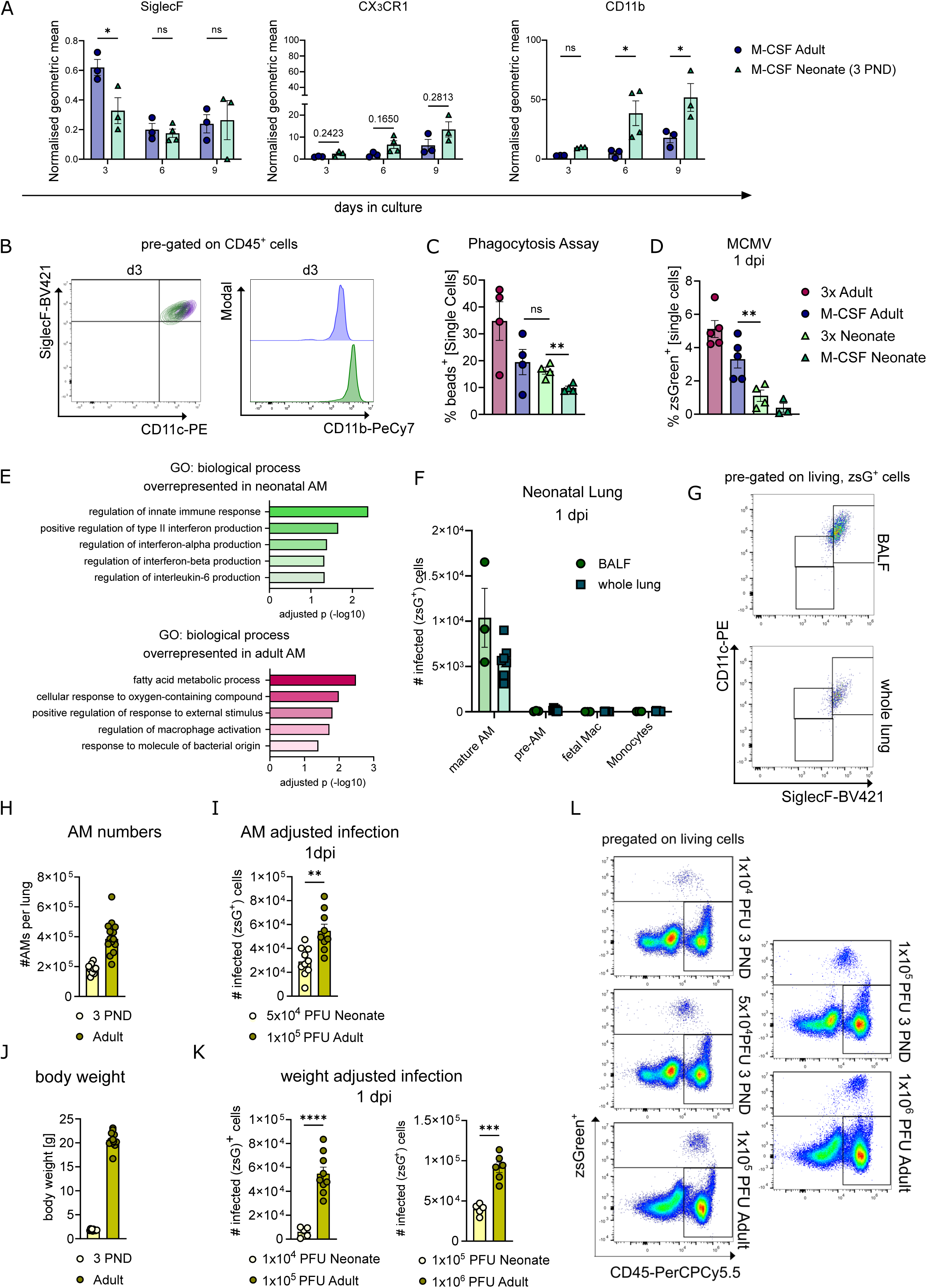
Neonatal mice are less susceptible to respiratory MCMV infections (A) Geometric mean of SiglecF, CX3CR1 and CD11b of M-CSF cultured adult and neonatal AM (isolated from 3 PND mice) normalised to the respective 3x cultured AM group across different timepoints and corresponding ages obtained by flow cytometry. *n=*3, 3 independent experiments, Tukey’s multiple comparisons test (B) Representative contour plot (left) comparing SiglecF and CD11c of 3x cultured (purple) and M-CSF cultured (dark green) AM of adult and 3 PND mice on day 3 (d3) of culture. Representative histograms (right) of CD11b on 3x cultured (purple) and M-CSF cultured AM (dark green) on d3 of culture. (C) Assessment of phagocytosis in 3x and M-CSF cultured AM with yellow-green fluorescent latex-beads using flow cytometry at d9 of culture of adult AM (purple, same data as in figure 3H) in comparison to neonatal AM (green). Data is depicted as % of single cells. *n=* 4, 3-4 independent experiments, unpaired t-test with Welch’s correction (D) Quantification of MCMV*^zsG^* (MOI 1) infected 3x and M-CSF cultured AM at 1 dpi of adult AM (purple, same data as in figure 3I) in comparison to neonatal AM (green). Data is depicted as % of single cells. *n=*3-5, 3-4 independent experiments, unpaired t-test with Welch’s correction (E) Gene ontology (GO) overrepresentation analysis of biological processes from bulk RNA-sequencing data (Figure 4) for top 500 differentially expressed genes (log2fc>1 and padj<0.05) neonatal AM (green) and adult AM (pink). (F) Absolute infected (zsGreen^+^) cell numbers in either whole lung (square shaped data points) or BALF (round shaped data points) of neonatal mice 1 dpi quantified by flow cytometry. Mice were infected with 1×10^5^ PFU *i.n.*, *n=*3 (BALF) and *n=*7 (whole lung), 1-2 independent experiments (G) Representative dot plots of infected cells 1 dpi in the neonatal BALF (top) and the whole lung (bottom), gated on SiglecF and CD11c. (H) AM numbers from 3 PND mice and adult mice, respectively. Data was obtained by FACS. *n=*10 (3 PND) and *n=*14 (adult) (I) Quantification of absolute infected (zsGreen^+^) cell numbers in either neonates (yellow) or adults (green) 1 dpi, assessed by flow cytometry. Mice were infected with 5×10^4^ PFU (neonates) or 1×10^5^ PFU (adults) *i.n., n=* 10 (neonate) and *n=* 9 (adult), 2 independent experiments, unpaired t-test with Welch’s correction (J) Body weight in gram (g) of 3 PND mice and adult mice, respectively. *n=*11 (3 PND) and *n=*12 (adult), 2 independent experiments (K) Quantification of absolute infected (zsGreen^+^) cell numbers in either neonates (yellow) or adults (green) 1 dpi, assessed by flow cytometry. Mice were infected with either 1×10^4^ PFU (neonates) and 1×10^5^ PFU (adults) *i.n.n=*5 (3 PND) and *n=*9 (adult) (left) or with 1×10^5^ PFU (neonates) and 1×10^6^ PFU (adults) *i.n. n=5* (3 PND) and *n=6* (adult) (right), 2 independent experiments, unpaired t-test with Welch’s correction. (L) Exemplary dot plots showing alive, MCMV*^zsG^* infected cells of neonates (top) and adults (bottom) 1 dpi. Data are represented as mean ± SEM. \**p <*0.05, \*\**p <*0.01, \*\*\**p <*0.001, \*\*\*\**p <*0.0001, *ns=* non-significant.

As outlined, CMV usually infects the respective host after birth, which does not lead to overt symptoms. Thus, we wondered whether lasting M-CSF imprinting in neonatal AM increased protection against MCMV. To test this, we infected neonatal mice at PND 3. Similarly to the situation in adult mice, only mature AM were infected 1 dpi, while pre-AM, fetal macrophages and monocytes were not infected in neither BALF nor in the lung tissue (Figure 5F, G). This was likely due to the exposed localization of mature AM in the alveoli, as analysis of BALF indicated their presence in the air-exposed space of the alveolus, while pre-AM, fetal mac and monocytes were not found in substantial numbers in the BALF (Figure S5A). Since AM represented the predominant and initial target of MCMV in both, adult (Figure 1) and neonatal mice (Figure 5F, G), we infected neonatal mice with an infectious dose, which was adjusted to the AM number, leading to an approximate halving of the adult dose per mouse (Figure 5H). With this approach, we found significantly less infected cells 1 dpi in neonates (Figure 5I, L). Next, we decided to confirm our findings additionally in a “weight adapted” model. Neonatal mice weigh approximately 10-times less than a young adult mouse (Figure 5J, S5B). Accordingly, we infected neonatal mice with 10-fold less PFU as compared to adult mice: either 10^4^ (neonate) vs 10^5^ (adult) PFU or 10^5^ (neonate) vs 10^6^ (adult) PFU. Again, they had less initially infected cells 1dpi (Figure 5K, L). It seems noteworthy that in this model, neonatal mice were infected even with higher PFU as compared to adult mice, when the infectious dose was normalized to the body weight (Figure S5B, C).

In summary, we found that neonatal AM retained a transient M-CSF-associated imprint that increased their plasticity and was associated with regulation of IFN responses and reduced susceptibility to MCMV infection. Consistently, neonatal mice displayed fewer infected cells following respiratory challenge.

## Discussion

Our study identifies AM rather than dendritic or epithelial cells as the indispensable initial niche for MCMV replication in the respiratory tract using a range of complementary pathogen-host fate mapping approaches. Moreover, it links their susceptibility to cytokine-driven AM state transitions shaped by M-CSF. Accordingly, AM represent an adjustable bottleneck for the establishment of a systemic viral infection.

Active human (H)CMV infection and viral replication in monocyte-derived macrophages is determined by base line levels of ISG (Schwartz et al., 2023). Our data suggest that AM susceptibility is highly dependent on their environmental input. M-CSF signalling induces a profound morphological, phenotypic and transcriptional reprogramming of AM. This transition is characterized by acquisition of MafB-associated gene expression, reduced phagocytic capacity, and enhanced IFN stimulated genes already in steady state. Functionally, this state confers reduced permissiveness to MCMV infection *ex vivo*, indicating that M-CSF provides environmental cues and in part determines an antiviral threshold in AM. It is an intriguing observation that M-CSF levels in the BALF increase after intranasal infection, thereby conceptually reverting the developmental trajectory from the lung airspace being a GM-CSF exclusive site.

*Ex vivo* culture is known to alter the phenotype of tissue-resident macrophages (Aktories et al., 2022; Subramanian et al., 2022), which is also to some degree reflected in our data (,e.g. *Vsig8* and *Fabp7*). However, by optimizing the culture conditions, limiting the culture duration as compared to previous studies and comparing conditions within our *ex vivo* system (3x versus M-CSF), we aimed to minimize these effects and observe M-CSF- and not culturing-specific effects.

The expression of the M-CSF receptor is largely restricted to the myeloid lineage during embryogenesis, with additional expression in placental trophoblasts (Hume et al., 2020). Although AM ultimately depend on GM-CSF for their differentiation and identity, they retain *Csf1r* expression, as indicated by reporter mouse models (Hawley et al., 2018) and confirmed here at the transcript level. However, the functional relevance of M-CSF receptor signalling in mature AM remained unclear.

In our study, we observed a reduction in AM numbers already in steady state in two mouse lines where *Csf1r* expression was reduced either in a macrophage-specific or non-selective fashion. In contrast, previous work reported comparable AM frequencies in >10-week-old *Csf1r*^ΔFIRE/ΔFIRE^ mice (Rojo et al., 2019). This discrepancy raises the question of whether the reduced AM compartment reflects impaired seeding from embryonic precursors, which might recover to normal AM numbers during adulthood, or an ongoing requirement for M-CSF receptor signalling in adult AM maintenance. Indeed, while yolk sac-derived macrophage progenitors critically depend on M-CSF (Gomez Perdiguero et al., 2015), fetal monocytes as defined by CD11b^hi^ F4/80^lo^ expression at E15.5, appear less sensitive to M-CSF receptor deficiency, at least in the skin (Hoeffel et al., 2012). In our dataset, fetal macrophages and early postnatal AM progenitors, *i.e.* pre-AM, exhibited a strong M-CSF signature that rapidly diminished as cells slowly mature into GM-CSF-dependent AM. Cross-species analyses supported a conserved transition from M-CSF-dominated to GM-CSF-dominated transcriptional programs during AM differentiation. Nonetheless, the M-CSF transcriptomic signature persisted in neonatal AM until PND 3. This developmental imprint had functional consequences: neonatal AM retained features of M-CSF exposure, including heightened plasticity and an antiviral transcriptional polarisation. This translated into reduced AM susceptibility to MCMV with neonatal mice exhibiting fewer infected cells than adult mice after respiratory infection.

Together, our findings support a model in which MCMV exploits a temporally and environmentally defined macrophage niche for early replication, while the intrinsic state of AM - shaped by cytokine signalling and developmental history - determines infection efficiency. Accordingly, this work extends the role of M-CSF well beyond its known function in macrophage maintenance. It positions M-CSF as a regulator of AM plasticity and antiviral readiness, and thus as a factor in the fine-tuning of early life host-pathogen interaction. More broadly, these findings illustrate how the ontogeny of tissue-resident macrophages and signals from the microenvironment interact to control viral tropism and may inform strategies to modulate susceptibility to respiratory infections.

## Acknowledgments

We are indebted to Adriana Greco, Reem Alsumati, Dominique Gütle and the Lighthouse facility (Freiburg) for their assistance with cell sorting and confocal microscopy, to Marco Prinz (Freiburg) and Steffen Jung (Israel) for provision of mice, and to the Center for Experimental Models and Transgenic Service (Freiburg) for excellent technical support. We also thank the Core Facility for Electron Microscopy (EMcore).

Z.R. received funding by the DFG TRR359 (Project ID 491676693). K.K. was supported by the DFG by project grants within the TRR359 (Project ID 491676693), TRR167 (Project ID 259373024), CRC1479 (Project ID 441891347), CRC1160 (Project ID 256073931) FOR5775 (Project ID 533863915), by the DFG under Germany’s Excellence Strategy (grant no. CIBSS—EXC-2189, Project ID 390939984) and by the Heisenberg program of the DFG (Project ID 544402801). J.K. received funding by the DFG TRR359 (Project ID 491676693). Sagar received funding by the DFG TRR359 (Project ID 491676693). P.H. received funding by the DFG (HE3127/9, HE3127/12, HE3127/16), TRR359 (Project ID 491676693), TRR167 (Project ID 259373024) and CRC1160 (Project ID 256073931). S.B. received funding from the DFG TRR359 (LAUNCH1) and together with J.K. was funded by the Hans A. Krebs Medical Scientist Programme (Faculty of Medicine, University of Freiburg). The Lighthouse Core Facility is funded in part by the Medical Faculty, University of Freiburg (Project Numbers 2023/A2-Fol; 2021/B3-Fol) and the DFG (Project Number 450392965). The Core Facility for Electron Microscopy (EMcore) at the University Freiburg Medical Center—IMITATE is registered with the DFG (German Research Foundation) under the reference number RI_00555.

## Methods

### Viruses

#### Cytomegaloviruses

Rescue of MCMV-GFP and MCMV-floxGFP from the bacterial artificial chromosomes (BACs) pSM3fr-MCK-2fl-M36GFP (Baasch et al. 2021) and pSM3fr-MCK-2fl-Dm157-flox-eGFP (Tegtmeyer et al., 2019), as well as pSM3fr-MCK-2fl-Dm157-hIE1zsGreen, was achieved by mouse embryonic fibroblasts (MEF) transfection. Subsequent propagation of all CMVs was performed on MEFs. Finally, purification of the viruses was conducted with a sucrose cushion.

#### Plaque assay for quantification of infectious MCMV particles

For quantification of infectious particles, a plaque assay was performed as previously described in (Reddehase et al., 1985). The assays were carried out in technical triplicates.

### Animal procedures

#### Mice

WT C57BL/6J mice were either purchased from Janvier Laboratories (Le Genest-Saint-Isle, France) or bred in the CEMT animal facility of the University Medical Centre Freiburg. All used transgenic mice (*Cx3cr1^gfp/+^, Cd169^Cre/+^; Csf1r^fl/fl^, Csf1r^ΔFIRE/ΔFIRE^, Csf2rb^-/-^, LysM^cre/+^, Itgax^cre/+^, Clec9a^cre/+^, Mrc1^creER/+^, Cx3cr1^creER/+^, S*ftpc*^creER/+^, CAG^creER/+^*) were bred in the aforementioned facility. Both genders were used for this study, as the examined criteria were sex-independent. Conducted animal experiments were approved by the Federal Ministry for Nature, Environment and Consumer’s protection of the federal republic Baden-Württemberg, Germany.

#### *In vivo* intranasal MCMV infection adult mice

For intranasal infection, 8–18-week-old C57B6/J mice were anesthetised with an intraperitoneal injection of Ketamin (100 mg/kg body weight) and Xylazin (10 mg/kg body weight). Afterwards, a total amount of 40 µL MCMV in PBS (1×10^5^ - 1× 10^6^ PFU) was administered intranasally. The mice were monitored until awakening and analysed at indicated time points.

#### *In vivo* intranasal infection neonatal mice

For intranasal infection of neonates, 3-day old C57B6/J mice were anesthetised with isoflurane (4% in O2). In the following, MCMV adjusted to 1×10^4^- 1×10^5^ PFU in 10 µL sterile PBS was administered intranasally. The mice were returned to the damp after the intervention.

#### AM depletion via clodronate-liposomes

For pharmaceutical depletion of AM with clodronate-liposomes, adult mice were anesthetised with an intraperitoneal injection of Ketamin (100 mg/kg body weight) and Xylazin (10 mg/kg body weight). Upon deep anesthesia, 40µL of 5mg/mL clodronate-liposomes (Clodrosome®, Encapsula Nano Sciences) were administered intranasally. Control-liposomes were installed accordingly.

### Extraction of bronchoalveolar lavage fluid (BALF)

To obtain BALF, the trachea was canulated with a catheter (BD Venlon^TM^ Pro). In the following, the lung was flushed multiple times with cold PBS and the BALF was collected on ice.

### Lung tissue preparation

For generation of lung single cell suspensions, mice were sacrificed using CO2. *Post mortem*, the heart was perfused with 5 mL of ice-cold PBS through the right ventricle. All lung lobes were excised, cut into small pieces and digested in PBS containing 10% FBS, Collagenase IV (250 µg/mL, Cat#: LS004188, Worthington), Hyaluronidase (1 mg/mL, Cat#: H3506-100MG, Sigma-Aldrich), Liberase^TM^ (20 µL/mL, Cat#: 5401119001, Roche) and DNase I (250 µg/mL, Cat#: DN25-1G, Sigma-Aldrich) using an environmental shaker for 60 min at 37°C and 180 rpm. For better distribution of the digestion mix the tubes were placed in a 45° angle. Following digestion, the suspension was vigorously vortexed, filtered through a 70 µm strainer and subsequently washed with cold FACS buffer (PBS, 1% FBS, 2 mM EDTA). Following centrifugation at 300 × g for 5 min, erythrocytes were lysed in 1 mL of cold eBioscience^TM^ 1x RBC lysis buffer (Cat#: 00-4333-57, Invitrogen) for 5 min at room temperature (RT). After an additional washing step and successive centrifugation, the cell suspension was resuspended in FACS buffer for further usage.

### Cell culture

#### Bone marrow-derived macrophages (BMDM)

To BMDM, adult mice were sacrificed and femurs and tibiae were harvested. Bones were rinsed with 60% isopropanol followed by sterile phosphate-buffered saline (PBS). Bones were then flushed using RPMI (supplemented with 10% fetal calf serum (FCS), 2% Ciprofloxacin (Fresenius Kabi)). Bone marrow cells were subsequently filtered through a 70 µm cell strainer. After centrifugation at 300 × g for 7 min, the cells were resuspended in supplemented RPMI containing 20 ng/mL recombinant M-CSF (Cat#: 315-02, Peprotech) and cultured in 10 cm^2^ Petri dishes. Medium was replaced on day 4. On day 6, BMDMs were enzymatically detached with accutase (Thermo Fisher) and re-seeded in supplemented RPMI.

#### Primary mouse AM culture

AM were FACS sorted according to their positivity for CD45, CD11c, SiglecF and CD206 into RPMI medium. For flow cytometry 75 000 cells were seeded into 48-well plates and cultured in RPMI medium (supplemented with 10 % FCS, 2% Ciprofloxacin (Fresenius Kabi)) containing either 20 ng/mL recombinant mouse GM-CSF (Cat#: 315-03, Peprotech), 10 ng/mL recombinant human TGF-β (Cat#: 100-21, Peprotech) and 1 µM Rosiglitazone (Cat#: R2408, Sigma-Aldrich) (= 3x) or 20 ng/mL recombinant mouse M-CSF (Cat#: 315-02, Peprotech). Medium supplemented with fresh (G)M-CSF, TGF- β and Rosiglitazone, respectively, was exchanged on day 4 and day 7 of culture. For analysis, AM were detached using accutase (Thermo Fisher) for 7 min at 37°C and additional cell scraping.

#### *In vitro* MCMV infection

For *in vitro* infection of AM and BMDM with MCMV*^zsG^* or MCMV*^cre^*, respectively, the cells were supplemented with medium containing the corresponding amount of virus for the indicated MOI. Subsequently, they were centrifuged with 450 × g for 30 min at 20 °C. After that, AM were incubated for 60 min in an incubator at 37°C. Lastly, AM were washed with sterile PBS two times and fresh medium was added.

#### *In vitro* phagocytosis assay

For assessment of phagocytic capacity *in vitro*, fluorescent yellow-green latex beads (1.0 µm mean particle size, Cat#: L1030-1ML, Sigma-Aldrich) were pre-opsonised in FBS (1:5) for 1 h at 37°C. Then, 1% (v/v) of the pre-opsonised beads diluted in RPMI-medium were added to the cells and incubated for 45 min at 37°C. After that, AM were washed with PBS twice and harvested using accutase and a cell scraper as described above.

### Flow cytometry and fluorescence-activated cell sorting (FACS)

#### Surface staining

Blocking of FcγII/III receptors on the cell surface was achieved by incubation of the samples with Fc-block (anti-CD16/32 antibody, Clone 93, BioLegend) in FACS buffer (PBS, 1%FCS, 2mM EDTA) for 10 min at 4°C. After that, the samples were stained with an antibody mastermix in FACS buffer for additional 20 min. Lastly, the cells were pelleted at 300 × g for 5 min and either resuspended in FACS buffer for flow cytometry/FACS or the samples were processed further.

#### Life/dead staining

To distinguish living cells from dead ones a fixable viability dye (eFluor^TM^780 Cat#: 65-0865-14, Invitrogen) diluted in PBS was applied to the samples following surface staining. After an incubation of 20 min, cells were washed with FACS buffer and pelleted at 300 × g for 5 min.

For subsequent flow cytometric analysis, samples were acquired with the 3-laser Gallios (Beckmann Coulter) or 4-laser cytometer Cytoflex S (Beckmann Coulter). FACS was performed using the MoFlo Astrios (Beckmann Coulter).

#### Antibodies for flow cytometry and FACS

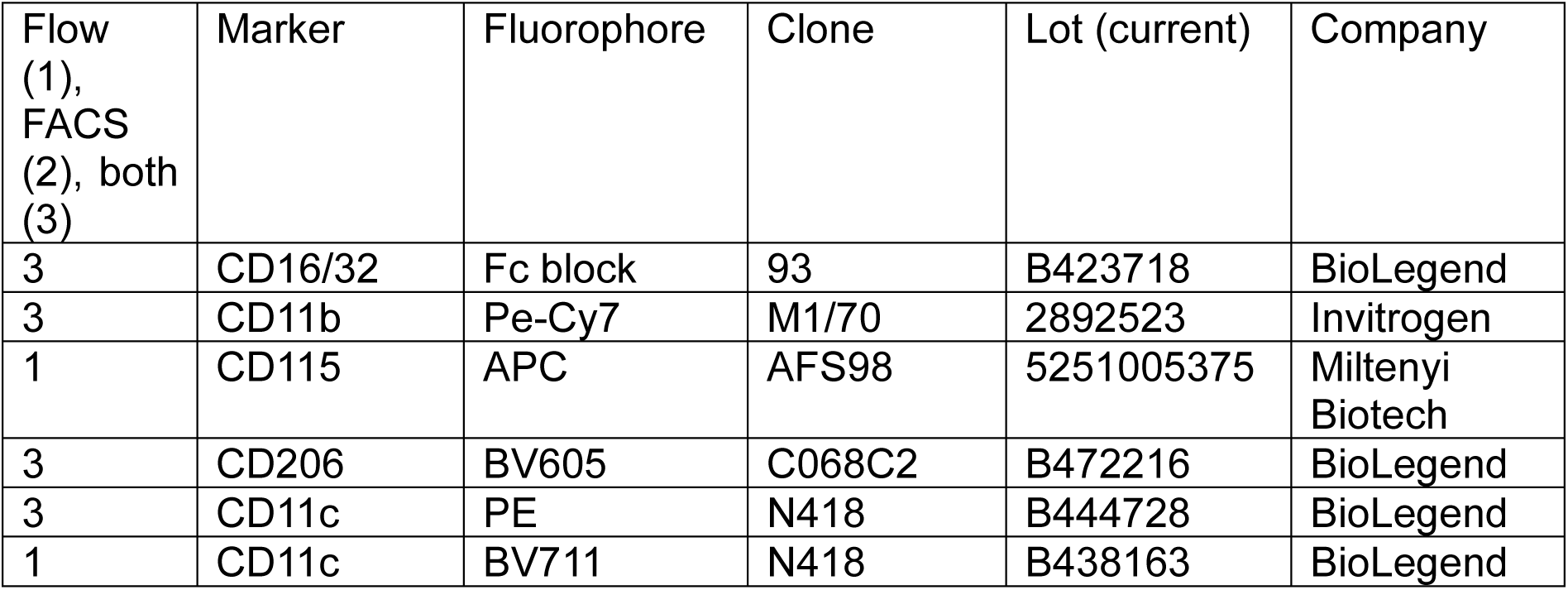
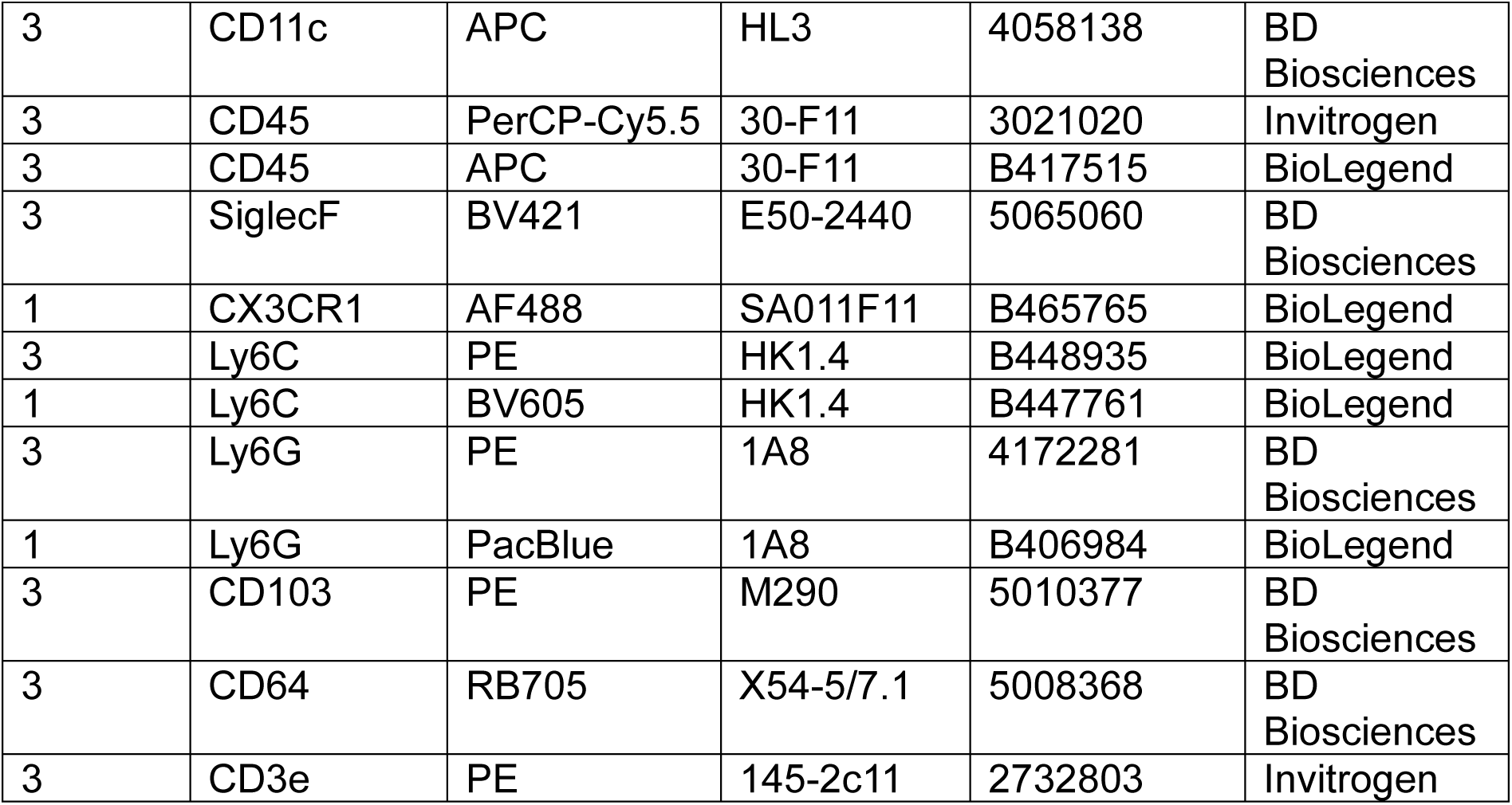

#### LEGENDplex^TM^ cytometric bead array

Prior to processing, BALF was cleared of cells and debris by centrifugation at 300 × g and 4°C for 5 min. For quantification of cytokines in the BALF, a LEGENDplex^TM^ array (Cat#:740622 and Cat#: 740677, BioLegend) was used according to the information provided by the manufacturer.

### Image techniques

#### Histology

For histology mice were sacrificed with CO2 and *post mortem* perfused with cold PBS through the right ventricle until the lungs were white. Subsequently, the trachea was canulated with a catheter and the lung was inflated with 4% PFA. Following 10-15 min of incubation at RT, the trachea was tied and the catheter removed. After that, the whole lung was excised and placed in 4% PFA at 4°C overnight. The next day, the lungs were washed with PBS once, transferred in 30% sucrose and placed at 4°c for dehydration. Three days later, the lungs were removed from sucrose and embedded in a cryomold containing OCT medium for snap-freezing on dry ice. For further processing, the lungs were cut in 20-40 µm thick slices using a cryotome and mounted on microscopy slides (Cat#: 473042491, TRAJAN Series 3 Adhesive).

#### Immunofluorescence staining lung

For staining, microscopy slides were permeabilised with 0.5 % TritonX-100 (Sigma-Aldrich) for 10 min at RT. Next, the slides were blocked with 10% goat-serum (Cat#: ab138478, Abcam) diluted in 0.05% TritonX-100 for at least 1h at RT. After 1 h, the blocking solution was discarded and the slides were incubated with primary/biotin-conjugated antibody at 4°C overnight. The next day, slides were washed at least three times and stained with secondary/conjugated/streptavidin-bound antibodies for 2 h at RT. For subsequent DAPI staining, the slides were incubated for 10 min at RT. Between the individual steps, the slides were washed 3-5 times. Lastly, the slides were mounted with Prolong diamond antifade mountant (Cat#: P36965, Thermo Fisher Scientific) and covered with a rectangular cover glass.

#### Immunofluorescence staining cells

AM were seeded into 8-well µ-slides (ibidi) and cultured as described above. Prior to staining, the cells were fixed in 4% PFA for 10 min at RT and then permeabilised for 5 min with 0.2% Tween-20 (Cat# P2287, Sigma-Aldrich). Cells were subsequently blocked with 0.05% Tween-20 containing 1% bovine serum albumin (BSA) (Cat#: A7906, Sigma-Aldrich) for 1 h at RT. In the following, cells were stained with primary antibody/biotin-conjugated antibody diluted in blocking solution overnight at 4°C. The next day, cells were stained with the secondary antibodies for 1-2h at RT and subsequently were subjected to DAPI staining for additional 10 min. Between the single steps, cells were washed with 0.05% Tween-20 at least twice. Finally, cells were mounted with fluorescence mounting medium (Cat#: S3023, Dako).

#### Scanning electron microscopy

AM were fixed in a mix of 2% glutaraldehyde and 0.1M cacodylate buffer (pH 7.4) at 4°C. Fixed cells were then dehydrated through a series of ascending ethanol concentrations for 15 minutes each (70%, 80%, 90%, 100%). After incubation in 50:50 ethanol/hexamethyldisilazane (Cat#: CAS-999-97-3; Sigma-Aldrich) and 100% hexamethyldisilazane, the solvent was allowed to evaporate. All samples were coated with platinum using an electron microscopy ACE600 Sputter-Coater (Leica). Samples were imaged using a Quattro <u>Scanning electron microscope</u> (Thermo Fisher Scientific).

### RNA-Seq and transcriptomic data analysis

#### RNA isolation

For RNA extraction, cells were lysed in RLT plus buffer containing 1%β-mercaptoethanol. In the following,RNA was extracted using the RNeasy Micro Kit according to manufacturer’s instructions (Cat#: 74004, QIAGEN). Samples were stored at-80°C until library preparation and further processing, performed by Novogene GmbH (Munich, Germany).

#### Bulk RNAseq of 3x AM, M-CSF AM, AM and interstitial macrophages (IM)

Bulk RNAseq was performed for AM and IM, as well as for 3x cultured AM and M-CSF cultured AM from adult *Cx3cr1^gfp/+^* reporter mice. For initial analysis steps the platform Galaxy Europe was used. Mapping of the data against the murine reference genome (*GRCm39*) was performed with STAR (Galaxy tool ID: toolshed.g2.bx.psu.edu/repos/iuc/rgrnastar/rna_star/2.7.11a+galaxy1). For generation of counting matrices, the tool featureCounts was used (Galaxy tool ID: toolshed.g2.bx.psu.edu/repos/iuc/featurecounts/featurecounts/2.0.8+galaxy0). The quality was subsequently assessed using the MultiQC tool (Galaxy tool ID: toolshed.g2.bx.psu.edu/repos/iuc/multiqc/multiqc/1.24.1+galaxy0). Next, in order to obtain normalised counts matrices of the individual samples, the DESeq2 package was used (Galaxy tool ID: toolshed.g2.bx.psu.edu/repos/iuc/deseq2/deseq2/2.11.40.8+galaxy0).

The further analysis was performed in R Studio. Volcano plots were generated using the ggplot2 R package. Differentially expressed genes were defined as abs(log2_FC)> 2 and *p* < 0.01. For heatmap generation the pheatmap tool was used. Before plotting, a log2-transformation was performed, followed by filtering out of all low expressed/not-expressed genes. Principal component analysis (PCA) was performed using the function prcomp with top 500 variable, log2 transformed normalised counts as input.

#### Extraction of AM and M-CSF signatures from bulk RNAseq of 3x AM, M-CSF AM, AM and IM

Differential expression analysis between conditions was performed using edgeR (Chen et al., 2016) (function calcNormFactors) and limma (Ritchie et al., 2015) (functions voom and lmFit), starting from the count matrix. Markers of a given condition were defined as genes with an adjusted p-value (BH method < 0.05) and log2FC> 1.

*Mafb* signature score (Vanneste et al., 2026) were calculated for each bulk RNA-sequencing sample as the mean DESeq2 variance-stabilized expression of detected genes in the *Mafb* gene set.

The AM and M-CSF signature was obtained from bulk RNA-sequencing data including freshly isolated AM and IM together with AM cultured with 3x or M-CSF.

First, markers were computed with the procedure previously described between AM vs IM. Then, markers were computed between 3x AM vs M-CSF AM. Finally, the intersection between markers of 3x AM and AM defined the AM signature, while the intersection between markers of IM and M-CSF AM defined the M-CSF signature.

Fisher’s test (R function fisher.test with default parameters) was computed between the 4 different clusters of genes previously defined in the human data and the M-CSF or AM signatures defined from the mouse bulk RNA-sequencing data. For this comparison, mouse gene names were converted to the corresponding human orthologs, when available, using the R library homologene. The background used in the Fisher test corresponds to all genes tested for differential expression analysis along the diffusion pseudo time. A cluster of genes was considered enriched for the M-CSF or AM signature if the Fisher test gave a p-value ≤ 0.001 and an odds ratio ≥ 2.

With these criteria, cluster 1 was the only one enriched for the AM signature, while clusters 3 and 4 were the only ones enriched for the M-CSF signature.

Applying the same criteria to the six clusters of genes differentially expressed along the trajectory in the mouse, clusters 5 and 6 were the only ones enriched for the AM signature, while cluster 4 was the only one enriched for the M-CSF signature. Relevant genes were shown on an heatmap generated with the r library pheatmap.

#### Bulk RNAseq of adult AM, neonatal AM fetal macrophages and pre–AM

Bulk RNAseq was performed for mature AM, fetal macrophages, and pre–AM of WT PND 3 mice, as well as for mature AM from WT adult mice, respectively.

Bulk RNAseq data were mapped with STAR (Dobin et al., 2013) (version 2.7.11b) on a genome index built with *Mus_musculus.GRCm39.dna.primary_assembly.fa* and *Mu s_musculus.GRCm39.111.gtf*. Count matrices for each sample were generated with featureCounts (Subread (Liao et al., 2013) version 2.1.1).

PCA embedding was computed using the R function prcomp (with scale. = TRUE) from the log-normalized count matrix (using the Seurat function NormalizeData (normalization.method = “LogNormalize”)), after keeping only genes expressed in at least 3 samples.

Differentially expressed genes were computed as previously described between adult AM and pre–AM.

Hierarchical clustering (function hclust with method = “ward.D2”) and cutreeHybrid (deepSplit = 0, minClusterSize = 50) were applied to the top 1k differentially expressed genes (obtained with the limma (Ritchie et al., 2015) function topTable with adjust = “BH”) between these two conditions,. This resulted in 3 clusters of genes (cluster 1: 433 genes; cluster 2: 420 genes; cluster 3: 147 genes)

#### Published human single cell RNA-Seq

In humans, to study the gene expression dynamics of AM from prenatal to postnatal stages, we used publicly available single-cell RNA-seq data (Wang *et al* 2020, Barnes *et al* 2024, Sikkema *et al* 2023).

For the prenatal stages (weeks 5–22 post-conception), cells annotated as APOE^+^ MΦ, CXCL9^+^ MΦ, and MΦ from (He et al., 2022), and APOE^+^ MΦ1, APOE^+^ MΦ2, CX3CR1^+^ MΦ, CXCL9^+^ MΦ, and SPP1^+^ MΦ from (Barnes et al., 2023) were merged together. After correcting for batch effects with the Seurat (Hao et al., 2021) function FindIntegrationAnchors (nfeatures = 2000), PCA embedding was computed with the Seurat function RunPCA (npcs = 30), and cluster analysis was performed with FindClusters (resolution = 0.1). This led to a total of 5 clusters.

Cluster 4 was subclustered (FindClusters, resolution = 0.1), since it was the only cluster mainly composed of APOE^+^ and CXCL9^+^ MΦ. Subclustering led to 2 subclusters, with one cluster strongly composed only of APOE^+^ MΦ (188 cells). This subcluster was kept for integration with postnatal stages of alveolar macrophages, since there is evidence showing APOE^+^ macrophages as precursors of AM.

For the postnatal stages, cells annotated as AM were selected from (Sikkema et al., 2022) at the one-month and six-month stages (46 cells), and from (Wang et al., 2020) at the 31-week and 3-year stages (4973 cells).

All postnatal AM and the prenatal APOE^+^ MΦ were integrated into a common count matrix. Batch correction was performed with the function RunHarmony (Korsunsky et al., 2019) (group.by.vars = “donor”, reduction = “pca”, theta = 0, lambda = 2) on the first 20 PCA components. The donor variable reflects the donor for the postnatal alveolar macrophages, while for the APOE^+^ MΦ it corresponds to the original paper. Afterward, cluster analysis was performed on the Harmony-corrected PCA components (FindNeighbors and FindClusters with resolution 0.1), leading to 4 clusters.

Trajectory was built using the R function slingshot (Street et al., 2018), using the Harmony-corrected PCA space as embedding and the clusters identified by Seurat after batch correction with Harmony. Pseudotime was assigned to each cell using the slingshot function slingPseudotime.

Spearman correlation was computed between pseudotime and real time points (from the fetal stage to 3 years of age) using the R function *cor.test*, yielding a correlation coefficient of 0.47 (p-value < 2.2 × 10⁻¹⁶).

To identify genes differentially expressed along the diffusion pseudotime, only genes with expression greater than 1 log-normalized count (using the Seurat function NormalizeData (normalization.method = “LogNormalize”)) in at least 100 cells were retained. Then, a GAM fit was performed on the log-normalized counts along the DPT using the R function mgcv (Wood, 2011) (with k = 3 and family = gaussian). For each gene, the Akaike Information Criterion (AIC) from the r library stats was computed for both the GAM fit and a constant model. A lower AIC balances goodness of fit with model complexity, and a lower value indicates a better fit.

The difference between the AIC of the constant fit and the AIC of the GAM fit was computed. Genes were sorted according to decreasing values of this difference (delta AIC). Only the top 1k genes with the highest delta AIC were retained for downstream analysis. Hierarchical clustering (function hclust with method = “ward.D2”) and cutreeHybrid (deepSplit = 0, minClusterSize = 50) were applied to the top 1k genes, using as input a distance matrix based on Spearman correlation computed from the GAM fitted values. Cluster analysis resulted in 4 clusters of genes (cluster 1: 414 genes; cluster 2: 228 genes; cluster 3: 185 genes; cluster 4: 173 genes). The mean trend per cluster was computed from the fitted values of each gene in that cluster.

GO terms for top 500 differentially expressed genes with padj < 0.05 and log2 fold change > 1, were determined with PANTHER (version 19.0, https://www.pantherdb.org/).

#### Published mouse single cell RNA-Seq

For studying the M-CSF signature during mouse development, AM were selected based on *Ear1*, *Ear2*, *Car4* expression from a publicly available mouse dataset (Cohen *et al*, 2018). This resulted in 4875 cells spanning time points from E12.5 to 40 hours postnatal.

scRNA-seq count matrices were processed and analyzed in R (v4.5.0) using the Seurat package (v5.3.0) (Hao et al., 2021). Low-quality cells were excluded based on the number of detected genes and the percentage of mitochondrial genes. The normalization method was set to ‘LogNormalize’. Dimensionality reduction was performed using the RunUMAP function and dims to 1:30. Default resolution was used for clustering. Cluster-specific genes were identified with the FindAllMarker function in Seurat with parameters logfc.threshold = 0.25. Trajectories were predicted using the Slingshot package(v 2.18.0) (Street et al., 2018), using the function slingshot with default settings and starting with the E12.5 developmental timepoint. The calculated trajectories were overlaid into the PCA embeddings. Diffusion pseudotime was assigned to each cell using slingPseudotime.

Spearman correlation was computed between pseudotime and real time points (from E12.5 to 40 hours postnatal) using the R function *cor.test*, yielding a correlation coefficient of 0.65 (p-value < 2.2 × 10⁻¹⁶).

To identify differentially expressed genes along the DPT, the same procedure previously described for human data was applied to the mouse data. This resulted in 6 clusters of genes (cluster 1: 311 genes; cluster 2: 181 genes; cluster 3: 177 genes; cluster 4: 158 genes; cluster 5: 118 genes; cluster 6: 55 genes).

### Software

Flow cytometric data were processed with either Kaluza (v.2.3, Beckman Coulter) or with FlowJo (v.10.10.0). Microscopy pictures were obtained with ZEN 3.0 SR (black) (v. 16.0.1.306) and subsequently processed using ZEN 3.0 (blue) (v. 3.0.79.0000). Graphical visualisation and statistical analysis was performed with GraphPad Prism (v.10.6.1.). For downstream analysis of bulk RNA-sequencing data, R Studio (2026.01.0 Build 392) was used.

## Statistical Analysis

For statistical analysis GraphPad Prism (v.10.6.1) was used. Statistical testing was performed as indicated in the figure legends.

---

**Supplemental figure 1.**
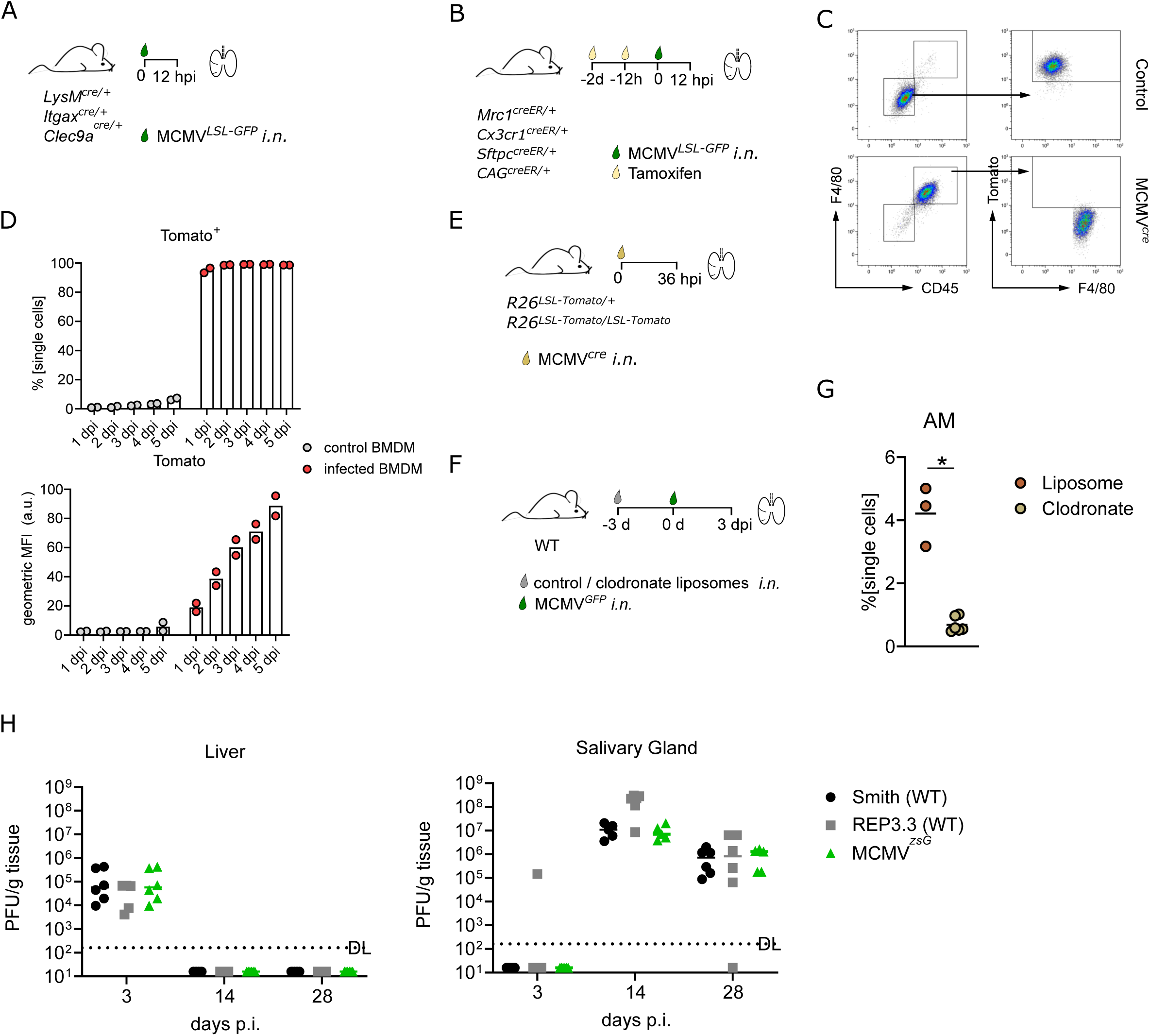
(A) Schematic of mice and MCMVs used for *in vivo* experiments Figure 1A-C. (B) Schematic of mice and MCMVs used for *in vivo* experiments Figure 1D-I. (C) Representative flow cytometry plots of BMDM from *R26^LSL-Tomato/+^* mice treated with PBS (top) or infected with MCMV*^cre^* (MOI5) at 2 dpi (bottom). (D) Quantification of Tomato^+^ infected BMDM (top) and of Tomato’s geometric mean fluorescence intensity (MFI, bottom) after infection of *R26^LSL-Tomato/+^* mice with MCMV*^cre^*. (E) Schematic of mice and MCMV used for *in vivo* experiments Figure 1J-L. (F) Schematic of mice, liposome administration and MCMV used for *in vivo* experiments Figure 1M, N. (G) Flow cytometric quantification of AM 3 days after administration of control liposomes or clodronate liposomes. Mann-Whitney test; \**p <*0.05 (I) BALB/c mice were infected intraperitoneally with 5×10^5^ PFU of either Smith strain of MCMV (Smith), the Mck2-repaired MCMV (Rep3.3) or repaired MCMV expressing zsG (MCMV*^zsG^*). PFU assays from liver and salivary gland were performed 3 dpi, 14 dpi and 28 dpi.

**Supplemental Figure 2:**
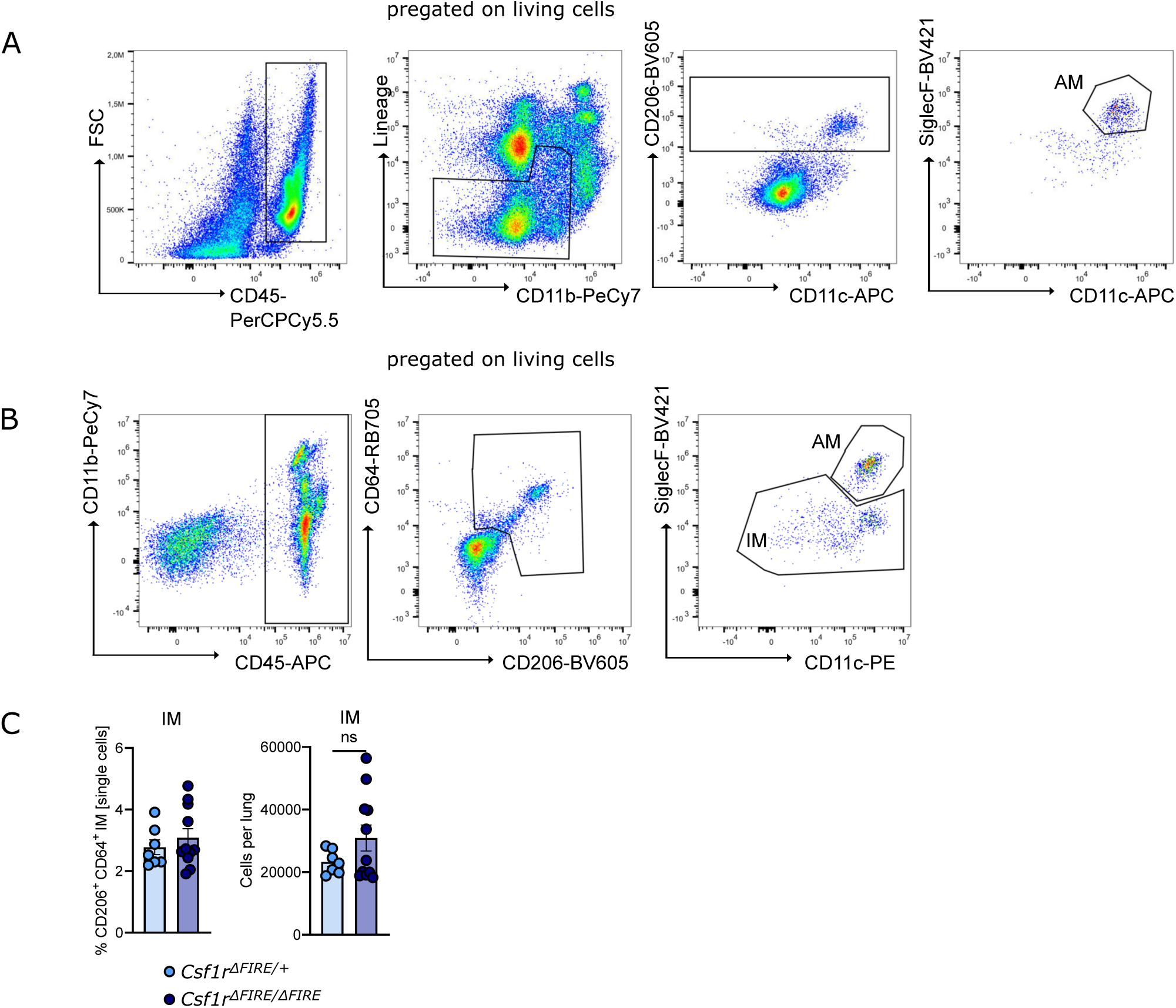
(A) Gating strategy for flow cytometric analysis of *Cd169^Cre/+^Csf1r^fl/fl^* mice. (B) Gating strategy for flow cytometric analysis of *Csf1r^ΔFIRE/ΔFIRE^* mice. (C) Quantification of absolute IM numbers and percentages in *Csf1r^ΔFIRE/ΔFIRE^* mice in comparison to *Csf1r^ΔFIRE/+^* mice assessed by flow cytometry. *n=*7-11, 4 independent experiments, unpaired t-test

**Supplemental Figure 3.**
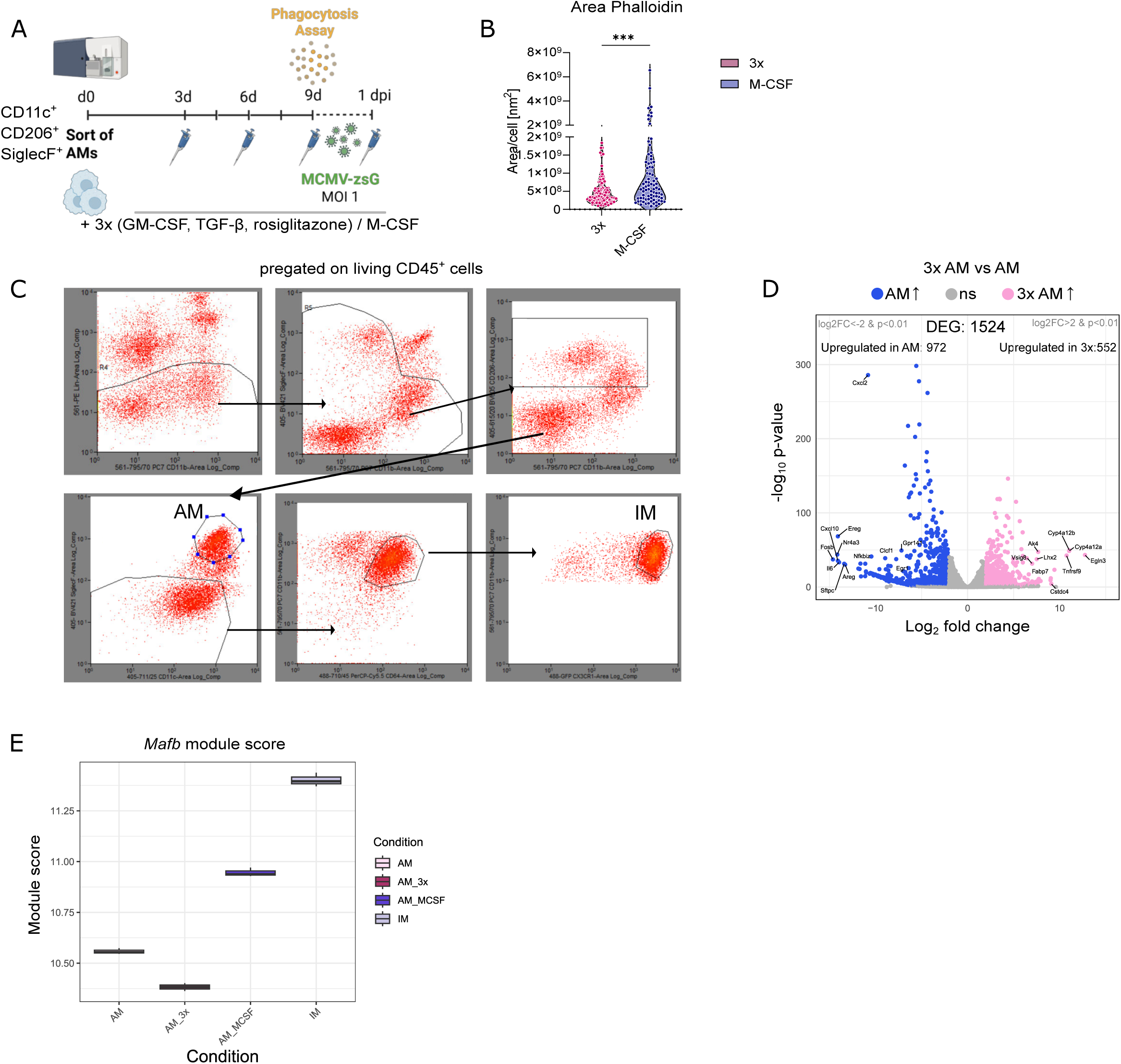
(A) Schematic illustration of workflow for primary mouse AM culture (created in biorender.com). (B) Quantification of cell area in nm^2^ of 3x and M-CSF culture AM based on 20 random cells counted per picture. Each dot is representative of one cell. *n=*3, unpaired t-test with Welch’s correction (C) Volcano plot showing differentially expressed genes (coloured dots: abs(log2_FC) > 2 and *p* < 0.01, labelled dots: abs(log2-FC) > 7, *p* < 0.01) that are upregulated in 3x AM (rosa) or in AM (purple) sorted from *Cx3cr1^gfp/+^* reporter mice. *n=*3 (D) Boxplots comparing *Mafb* module scores (Vanneste et al., 2026) between bulk RNA-sequenced 3x AM, M-CSF AM, AM and IM. n=3. *Mafb* signature scores were calculated per mouse as the mean variance-stabilized expression of genes in the *Mafb* gene set. Data are represented as mean ± SEM. \*\*\**p <* 0.001

**Supplemental Figure 4.**
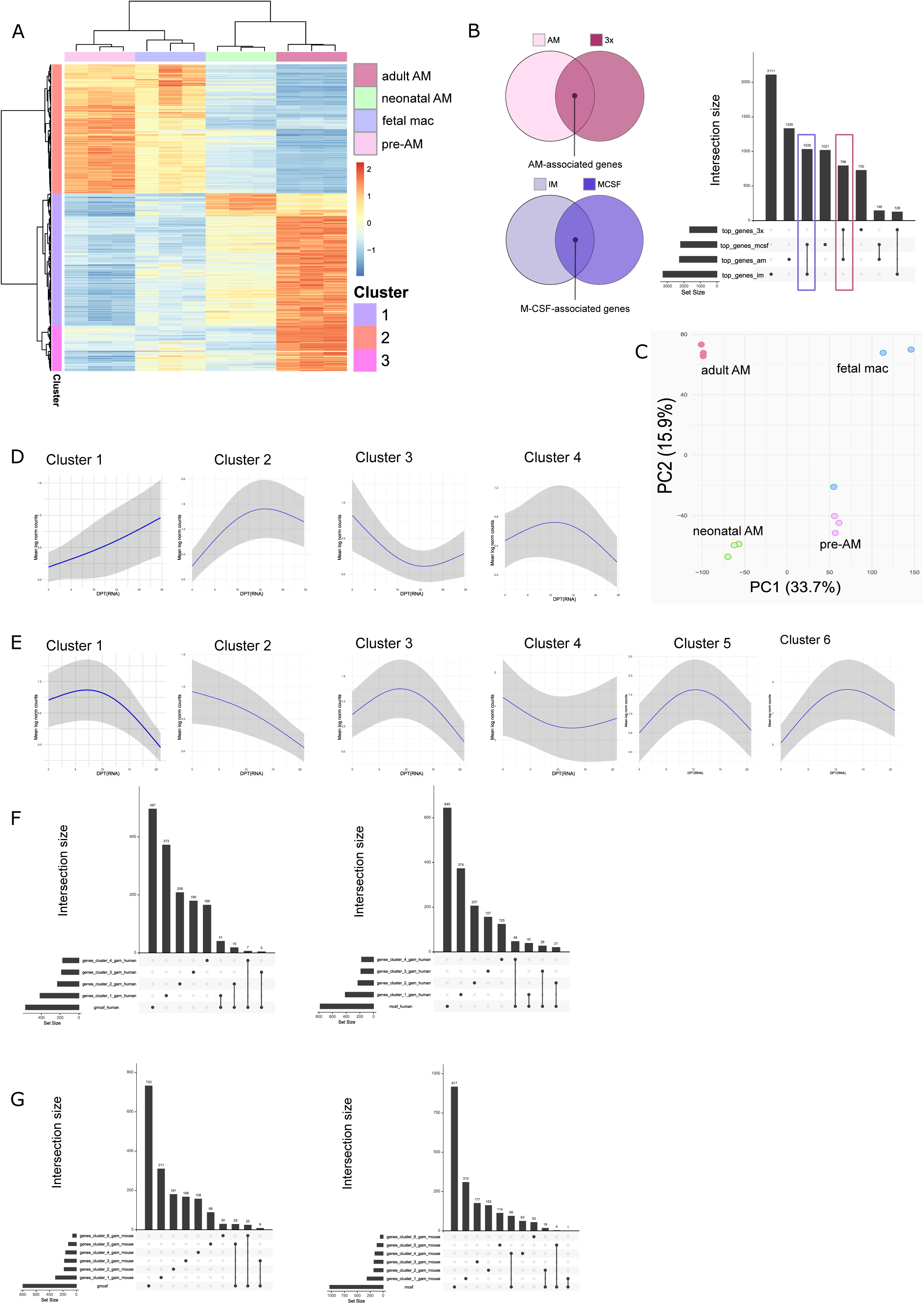
(A) Heatmap of top 1000 differentially expressed genes between adult AM and pre–AM, separated into 3 gene expression clusters based on increasing and decreasing gene expression across cell types. (B) Schematic illustration of how AM and M-CSF signatures were deducted from bulk RNA-sequencing data shown in Figure 3 (left) and upset plot showing the intersections between AM and 3x AM, as well as IM and M-CSF AM, respectively (right). (C) Average profiles obtained from the fitted values of genes belonging to cluster 1-4, which are composed of gene groups with specific trajectory patterns across pseudo time (human data). Mean profile +- Standard deviation (SD) is shown. (D) Average profiles obtained from the fitted values of genes belonging to cluster 1-6, which are composed of gene groups with specific trajectory patterns across pseudo time (mouse data). Mean profile +- SD is shown. (E) Upset plot showing the intersection between the cluster of differentially expressed genes along diffusion pseudo time defined in human and the M-CSF and AM signature. See methods for more details. (F) Upset plot showing the intersection between the cluster of differentially expressed genes along diffusion pseudo time defined in mouse and the M-CSF and AM signature. See methods for more details.

**Supplemental Figure 5:**
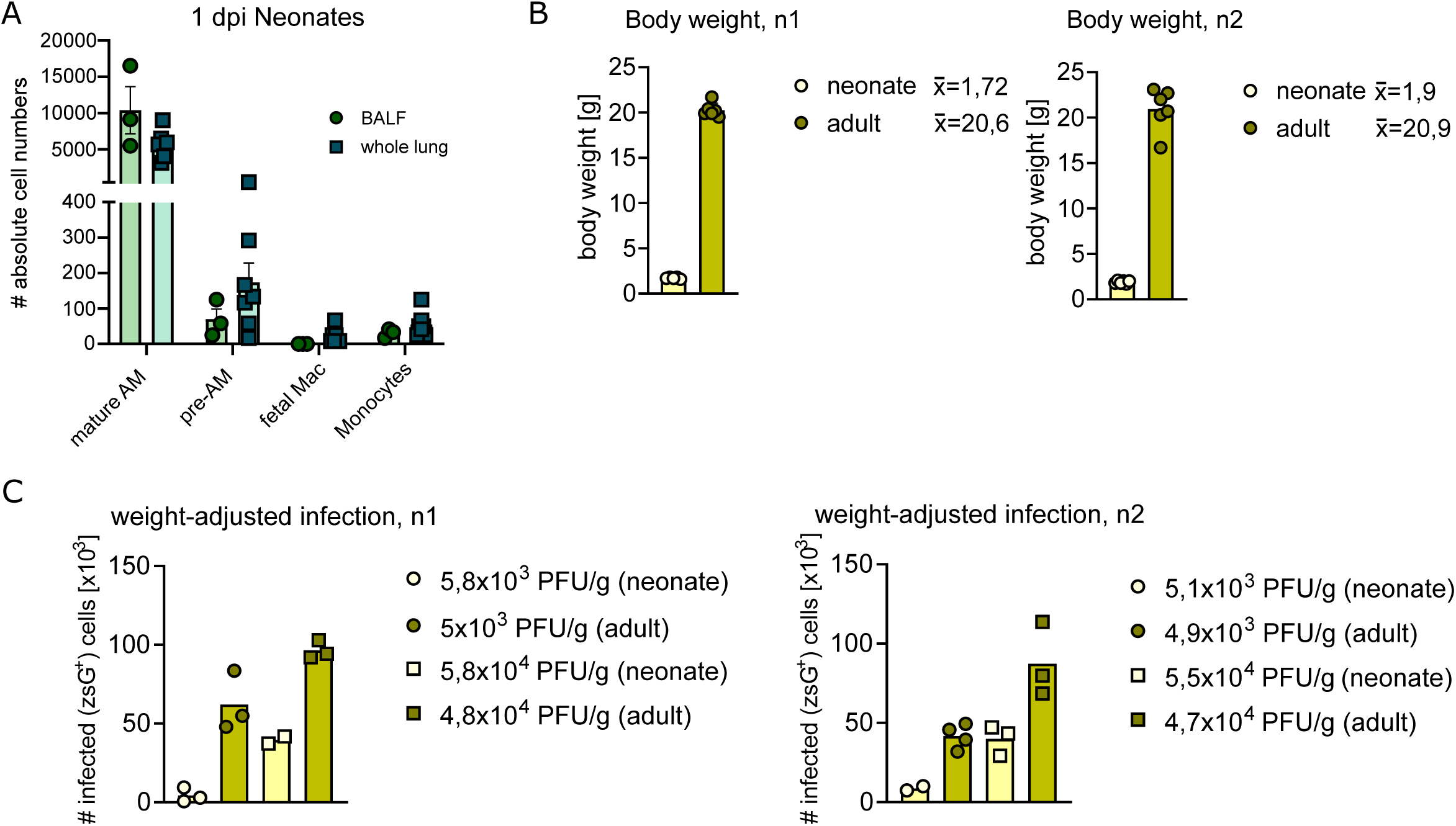
(A) Absolute cell numbers of neonatal mature AM, pre-AM, fetal mac and monocytes in either whole lung (square shaped data points) or BALF (round shaped data points) of neonatal mice 1 dpi quantified by flow cytometry. Mice were infected with MCMV*^zsG^* (1×10^5^ PFU *i.n.*), *n=*3 (BALF) and *n=*7 (whole lung), 1-2 independent experiments. (B) Body weight in gram (g) of PND 3 mice and adult mice, respectively, separated according to the two independent experiments. (C) Quantification of absolute infected (zsGreen^+^) cell numbers in either neonates (yellow) or adults (green) 1 dpi, assessed by flow cytometry and displayed as two different experiments (n1 and n2). Mice infected with either 1×10^4^ PFU (neonates, yellow circles) and 1×10^5^ PFU (adults, green circles) *i.n. n=*5 (3 PND) and *n=7* (adult) or with 1×10^5^ PFU (neonates, yellow squares) and 1×10^6^ PFU (adults, green squares) *i.n. n=5* (3 PND) and *n=6* (adult), the applied viral dosage was subsequently normalized to gram body weight.

